# STARCH SYNTHASE 4 is required for normal starch granule initiation in amyloplasts of wheat endosperm

**DOI:** 10.1101/2021.01.29.428798

**Authors:** Erica Hawkins, Jiawen Chen, Alexander Watson-Lazowski, Jennifer Ahn-Jarvis, J. Elaine Barclay, Brendan Fahy, Matthew Hartley, Frederick J. Warren, David Seung

**Affiliations:** John Innes Centre, Norwich Research Park, Norwich, NR4 7UH, UK; Quadram Institute, Norwich Research Park, Norwich, NR4 7UQ, UK

**Keywords:** amyloplast, BGC1, endosperm, granule initiation, SS4, starch, starch synthesis, wheat

## Abstract

- Starch granule initiation is poorly understood at the molecular level. The glucosyltransferase, STARCH SYNTHASE 4 (SS4), plays a central role in granule initiation in Arabidopsis leaves, but its function in cereal endosperms is unknown. We investigated the role of SS4 in wheat, which has a distinct spatiotemporal pattern of granule initiation during grain development.
- We generated TILLING mutants in tetraploid wheat (*Triticum turgidum*) that are defective in both SS4 homoeologs. The morphology of endosperm starch was examined in developing and mature grains.
- SS4 deficiency led to severe alterations in endosperm starch granule morphology. During early grain development, while the wild type initiated single ‘A-type’ granules per amyloplast, most amyloplasts in the mutant formed compound granules due to multiple initiations. This phenotype was similar to mutants deficient in B-GRANULE CONTENT 1 (BGC1). SS4 deficiency also reduced starch content in leaves and pollen grains.
- We propose that SS4 and BGC1 are required for the proper control of granule initiation during early grain development that leads to a single A-type granule per amyloplast. The absence of either protein results in a variable number of initiations per amyloplast and compound granule formation.

## INTRODUCTION

Starch is a major storage carbohydrate in leaves and non-photosynthetic organs of many plants. The starch-rich endosperm of cereal grains is an important source of calories in human diets. Starch forms insoluble semi-crystalline granules that are composed of the glucose polymers - amylopectin and amylose. The biosynthesis of these polymers is relatively well understood and conserved among different plants (Smith & Zeeman, 2020). By contrast, we are only beginning to understand the mechanism of starch granule initiation, and there is vast diversity in the number and morphology of granules between different species and organs (Seung & Smith, 2019).

There are five major classes of active starch synthases - SS1, SS2, SS3, SS4 and GBSS – which are glucosyltransferases that elongate α-1,4-linked glucan chains of starch polymers using ADP-glucose. SS1, SS2 and SS3 are involved in amylopectin synthesis, and mutants of Arabidopsis and cereals defective in these isoforms have altered amylopectin structure (Wang *et al*., 1993; Morell *et al*., 2003; Zhang *et al*., 2005, 2008; Delvallé *et al*., 2005; Fujita *et al*., 2006, 2007; Szydlowski *et al*., 2011). Amylopectin synthesis also requires branching enzymes (BEs) and debranching enzymes (isoamylases - ISAs) (Delatte *et al*., 2005; Dumez *et al*., 2006; Sundberg *et al*., 2013). Granule-Bound Starch Synthase (GBSS) is required for amylose synthesis (Seung, 2020). In Arabidopsis leaves, SS4 is required for both normal granule initiation and morphogenesis, but does not make a major contribution to amylopectin structure (Roldán *et al*., 2007; Szydlowski *et al*., 2009; Crumpton-Taylor *et al*., 2012, 2013; Seung *et al*., 2017; Lu *et al*., 2018). While chloroplasts of wild-type leaves contain multiple granules, those of the *ss4* mutant typically contain only one or no granule. The granules of *ss4* have distinct spherical morphology, rather than the flattened shape of wild-type starch granules. The *ss4* mutant also accumulates ADP-glucose, suggesting that other SS isoforms cannot effectively utilise this substrate in the absence of SS4 (Crumpton-Taylor *et al*., 2013; Ragel *et al*., 2013).

Arabidopsis SS4 acts at least partially in complex with other proteins that are required for normal granule initiation. These include PROTEIN TARGETING TO STARCH family members, PTST2 and PTST3 (Seung *et al*., 2017). PTST2 is proposed to play a role in delivering maltooligosaccharide primers to SS4 for further elongation (Seung *et al*., 2017). SS4 interacts with a coiled-coil protein, MRC (also called PII1), but the exact role of this interaction is unknown (Seung *et al*., 2018; Vandromme *et al*., 2019).

The function of SS4 in non-photosynthetic amyloplasts of storage organs and seeds is unknown. Granule initiation patterns in the endosperm of the Triticeae are radically different from those in Arabidopsis leaves: large, flattened A-type granules initiate early during grain development, and small round B-type granules initiate 10-15 days after the A-type granules (Bechtel *et al*., 1990; Howard *et al*., 2011; Chia *et al*., 2020). Nonetheless, the loss of PTST2 orthologs – FLOURY ENDOSPERM 6 (FLO6) in barley and B-GRANULE CONTENT 1 (BGC1) in wheat - has major effects on granule initiation in the endosperm (Suh *et al*., 2004; Saito *et al*., 2017; Chia *et al*., 2020). This discovery raises the possibility that granule initiation in wheat endosperm is via an SS4-containing complex, similar to that in Arabidopsis leaves. Here we aimed to generate and characterise wheat mutants that are deficient in *Ta*SS4, to determine its role in starch synthesis in the endosperm. The mutants had highly abnormal endosperm starch morphology, resulting from the formation of compound starch granules. Interestingly, this phenotype resembled mutants defective in *Ta*BGC1 (Chia *et al*., 2020). Our work demonstrates that both *Ta*SS4 and *Ta*BGC1 are required for the control of granule initiation in endosperm amyloplasts.

## MATERIALS AND METHODS

### Bioinformatics analyses

*Ta*SS4 loci (Fig. 1) were identified using BLAST against the wheat RefSeq 1.1 genome of cultivar Chinese Spring (Appels *et al*., 2018) on Ensembl plants (Kersey *et al*., 2018). *Ta*SS4 sequences from cultivars Cadenza, Claire, Kronos, Paragon and Robigus were obtained from the Grassroots database (Clavijo *et al*., 2017).

**Fig. 1.**
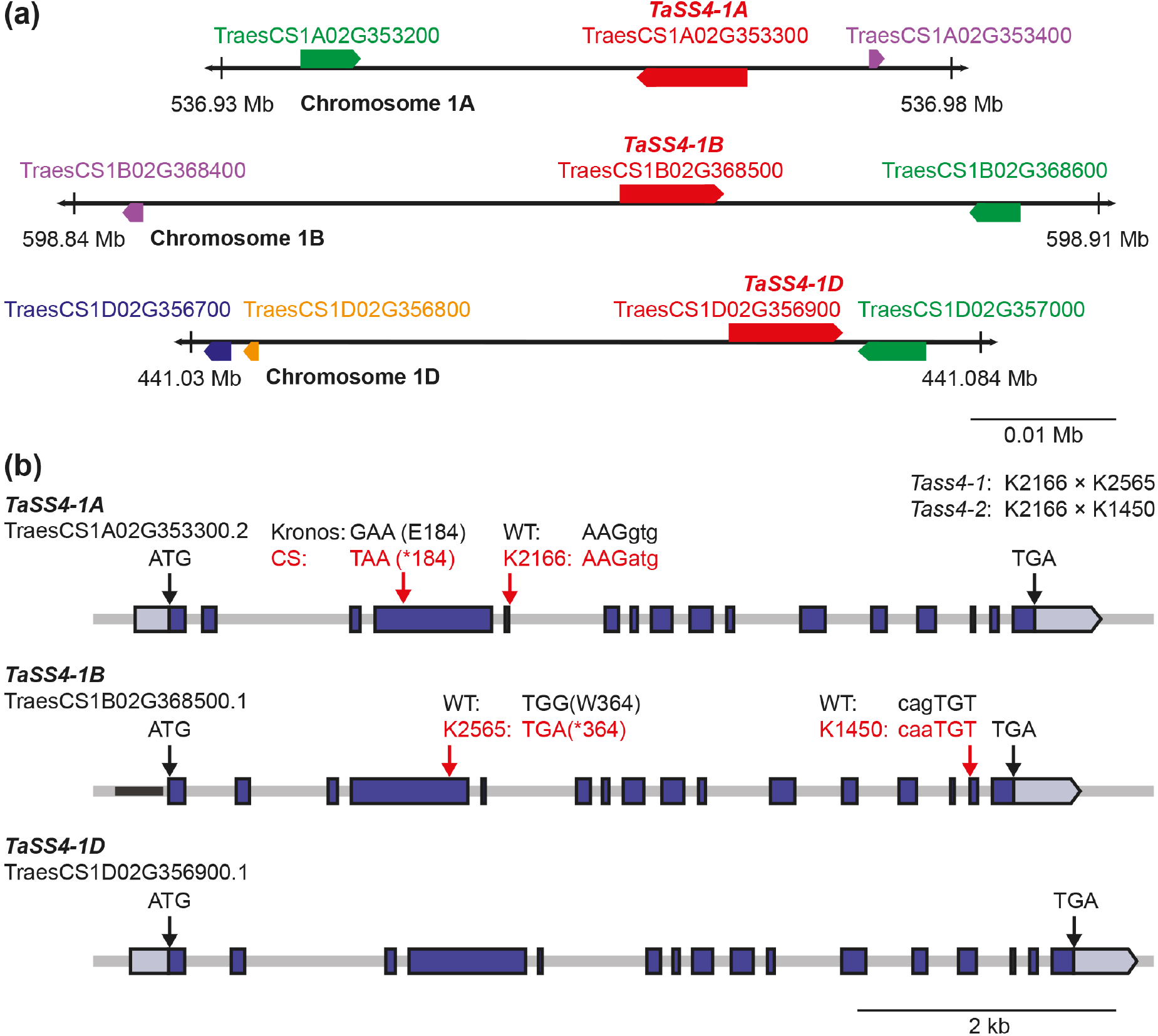
Schematic illustrations of *TaSS4* homoeologs. **(a)** Location of *TaSS4* homoeologs on group 1 chromosomes. The red boxes represent *TaSS4* homoeologs, while homoeologs of the adjacent genes are shown in green (phosphodiesterase-like protein), purple (glycoside hydrolase family 18 protein), blue (P loop-containing nucleoside triphosphate hydrolase) and orange (serine acetyltransferase-like protein). Arrowheads on the boxes indicate direction of transcription. Chromosome coordinates are indicated below each region. **(b)** Gene models of the *TaSS4* homoeologs. Exons are represented with blue boxes, while light blue boxes represent the 5′ and 3′ UTRs. Mutations in the *Tass4-1* and *Tass4-2* mutant lines are depicted with red arrows and the mutated codons/amino acids are shown in red letters. The polymorphism in *TaSS4-1A* between Kronos and Chinese Spring (CS) is indicated.

*Ta*SS4 and *Ta*BGC1 transcript levels during tetraploid wheat grain development were extracted from the datasets of Maccaferri *et al*. (2019) and Xiang *et al*. (2019). Raw RNA-Seq reads obtained from the GenBank Sequence Read Archive (SRA) were processed using Trimmomatic (Bolger *et al*., 2014) to remove adapter sequences. Processed reads were aligned to the *Triticum turgidum* transcriptome (Maccaferri *et al*., 2019) using the Quasi align mode in Salmon (Patro *et al*., 2017) outputting normalised expression as transcripts per million (TPM). Transcript levels of *Ta*SS4 in different organs of hexaploid wheat were retrieved from the wheat expression database (http://www.wheat-expression.com)(Borrill *et al*., 2016).

### Plant materials and growth

Mutants in *Triticum turgidum* (cultivar Kronos) were identified using the wheat *in silico* TILLING resource (http://www.wheat-tilling.com)(Krasileva *et al*., 2017): Kronos2166(K2166) for *TaSS4-1A*, Kronos2565(K2565) and Kronos1450(K1450) for *TaSS4-1B*, and Kronos2275(K2275) for *TaBGC1-4B. TaBGC1-4A* mutants, Kronos2244(K2244) and Kronos3145(K3145) are from Chia *et al*. (2020). Plants were crossed to combine A- and B-homoeolog mutant alleles. AA BB, *aa* BB, AA *bb* and *aa bb* genotypes were selected in the F2 generation using KASP V4.0 genotyping (LGC) with the primers in Table S1.

Wheat plants were grown in controlled environment rooms (CER) or glasshouses at 60% relative humidity with 16 h light at 20°C and 8 h dark at 16°C. The CER light intensity was 300 μmol photons m^−2^ s^−1^. Experiments on leaves and developing grains were carried out on CER-grown material, whereas experiments with mature grains were carried out on either CER or glasshouse-grown material. *Nicotiana benthamiana* plants were grown in glasshouses set to provide a minimum of 16 h light at 22°C, and a dark period of 20°C. *Arabidopsis thaliana* plants were grown in CERs at 60% relative humidity, 12 h light (150 μmol photons m^−2^ s^−1^)/12 h dark cycles and constant temperature of 20°C.

### Starch purification, granule morphology and size distribution

Endosperm starch purification: Mature grains were soaked overnight in ddH_2_O at 4°C, then homogenised in a mortar and pestle with excess ddH_2_O. Developing grains (stored at −80°C post-harvest) were thawed prior to endosperm dissection and immediately homogenised in ddH_2_O using a ball mill at 30 Hz for 1 min. The homogenates were filtered (70 μm nylon mesh) then centrifuged, and the pellet was resuspended into 90% (*v/v*) Percoll, 50 mM Tris-HCl, pH 8 and centrifuged at 2500*g*, 5 min. The pellet was washed twice in 50 mM Tris-HCl, pH 6.8, 10 mM EDTA, 4% SDS (*v/v*), 10 mM DTT and resuspended in ddH_2_O.

Granule morphology was observed using a Nova NanoSEM 450 (FEI) scanning electron microscope (SEM). For cross-polarised light microscopy, the granules were imaged with a DM6000 microscope fitted with a DFC 320F camera (Leica). For analysis of granule size distributions, the starch was suspended into Isoton II (Beckman Coulter), and relative volume vs. diameter plots were generated using a Multisizer 4e Coulter counter (Beckman Coulter) with a 70 μm aperture tube. A minimum of 100,000 particles was measured per sample. All measurements were conducted with logarithmic bin spacing but are presented on a linear x-axis for clarity. The mean diameters of A- and B-type granules, and relative volume fraction of B-type granules, were calculated by fitting a mixture of two log-normal distributions in R (script available at https://github.com/JIC-CSB/coulter_counter_fitting).

### Starch quantification, composition and amylopectin structure

Grain starch quantification: Flour (milled in a ball mill; 5-10 mg) was dispersed in 20 μL 80% EtOH, and then incubated with 500 μL thermostable α-amylase in 100 mM sodium acetate buffer, pH 5, at 99°C for 7 min. Amyloglucosidase was added and incubated at 50°C for 35 min. All enzymes and reagents were from the Total Starch Assay kit (Megazyme, K-TSTA). The sample was centrifuged at 20,000*g* for 10 min. Glucose was measured in the supernatant using the hexokinase/glucose-6-phosphate dehydrogenase assay (Roche), for calculation of starch content in glucose equivalents. Leaf starch quantification: 10-day-old seedlings were harvested at the base of the lowest leaf and flash frozen in liquid N_2_. The material was homogenised in 0.7 M perchloric acid using a ball mill at 30 Hz. Insoluble material was pelleted by centrifugation, washed three times in 80% ethanol, then resuspended in water. Starch was digested using α-amylase/amyloglucosidase (Roche), and glucose was assayed as for grains.

Starch chain length distribution: Purified starch was solubilised and enzymatically debranched using methods adapted from Wu *et al*. (2014), and analysed using high performance size exclusion chromatography (HPLC-SEC) as detailed in Tuncel *et al*. (2019). Calibration curves were generated using pullulan standards (PSS-pulkit, Polymer Standard Service) having peak molecular weights ranging from 342 to 708,000 Da and with correlation coefficients of R^2^ = 0.9997 ± 0.0002. The calibration curves were used to determine the relationship between elution volume and hydrodynamic radius (V_h_) for the linear glucans, as described by Cave *et al*. (2009). The refractive index elution profiles were converted to SEC weight distributions as described by Perez-Moral et al. (2018). Amylose content was determined from chain length distributions as described by Vilaplana *et al*., (2012). Briefly, the cut-off between amylose and amylopectin in the chain length distribution was set at 100 degrees of polymerisation (D.P.), and the peak areas of the amylopectin (chains <100 D.P.) and amylose (chains >100 D.P.) were integrated. Amylose content was estimated as the ratio of the amylopectin and amylose peak areas expressed as a percentage.

### Light and transmission electron microscopy of sections

Mature grain sections: After transverse grain bisection with a razor blade, thin 1 μm sections were produced from the cut surface using an Ultracut UC6 microtome (Leica) fitted with a glass knife. Sections were stained with a 1 in 20 dilution of Lugol’s iodine solution (Sigma), and mounted in Histomount (National Diagnostics). Light microscopy was carried out on an AxioObserver Z1 microscope with an AxioCam camera (Zeiss); or a DM6000 microscope with a DFC 320F camera (Leica).

Leaf/developing grain sections: Leaf segments were excised from approximately halfway along the length of a flag leaf (for wheat) or a young rosette leaf (for Arabidopsis), and fixed in 2.5% *(v/v)* glutaraldehyde in 0.05 M sodium cacodylate, pH 7.3 at 4°C. Developing grains (15 days post anthesis - dpa) were cut in half before immersion in fixative. Using an EM TP embedding machine (Leica, Milton Keynes, UK), samples were post-fixed in 1% *(w/v)* OsO_4_ in 0.05 M sodium cacodylate for two hours at room temperature, dehydrated in ethanol and infiltrated with LR White resin (London Resin Company). LR White blocks were polymerised at 60°C for 16 h. For light microscopy, the semi-thin sections (0.5 μm) were prepared. Leaf sections were stained with reagents from the Periodic Acid Schiff kit (Abcam, ab150680), by incubating 30 min in the periodic acid solution and 5 min in Schiff’s reagent, then staining with 1% *(w/v)* toluidine blue for 30 sec prior to mounting in Histomount. Sections from developing grains were stained with 1% *(w/v)* toluidine blue. Light microscopy was carried out as described above. For transmission electron microscopy (TEM), ultrathin sections (~80 nm) were cut with a diamond knife and placed on formvar and carbon-coated copper grids (EM Resolutions). The sections were stained with 2% *(w/v)* uranyl acetate for 1 h and 1% *(w/v)* lead citrate for 1 min, washed in distilled water and air dried. Sections were viewed on a Talos 200C TEM (FEI) at 200 kV and imaged with a OneView 4K x 4K camera (Gatan).

### Visualisation and scoring of starch in pollen

Mature anthers were harvested into 80% *(v/v)* EtOH and stained with a 1 in 20 dilution of Lugol’s iodine solution (Sigma) overnight. After destaining in ddH_2_O, pollen was observed with light microscopy as described above. The percentage of starchless pollen (no visible iodine stain) was calculated by scoring the first ≈100 pollen grains observed.

### Cloning and transformation of plant material

*Ta*SS4-1A, *Ta*SS4-1B, *Ta*BGC1-4A and *Ta*BGC1-4B coding sequences were codon optimised to ease sequence complexity and synthesised as gBlocks gene fragments (IDT DNA), flanked with attB1 and attB2 Gateway recombination sites. The optimised sequences are provided in Table S2. The fragment was recombined into the pDONR221 vector using BP Clonase II (Thermo-Fisher). The sequences were recombined into pUBC-YFP (Ubiquitin10-driven expression and C-terminal YFP-tag)(Grefen *et al*., 2010) or pJCV52 (CaMV 35S-driven expression and C-terminal HA-tag)(Karimi *et al*., 2002).

For transient expression in *Nicotiana benthamiana, Agrobacterium tumefaciens* (strain AGL-1 or GV3101) harbouring the relevant constructs were grown at 28°C for 48 h. Cultures were resuspended in ddH_2_O at OD_600_ = 1.0, and infiltrated into the abaxial leaf surface using a syringe. Proteins were extracted 48-72 h after infiltration. The *TaSS4* 1B-YFP:pUBC-YFP construct was transformed into Arabidopsis by floral dipping (Zhang *et al*., 2006).

Transformants were selected in the T_1_ generation using the Basta resistance marker. Basta-resistant individuals from the T_2_ or T_3_ generation (heterozygous or homozygous for the transgene; single or multiple insertions) with *Ta*SS4 expression confirmed using immunoblots were used for experiments.

### Production of antibodies and immunoblotting

To produce *Ta*SS4 and *Ta*BGC1 antibodies, the coding sequence of the proteins (minus transit peptide) were amplified using primers in Table S1, and *Ta*SS4-1B:pDONR221 or *Ta*BGC1-4B:pDONR221 as templates. The amplicons were cloned into the pProExHTb vector (Invitrogen) in frame with the N-terminal His_6_-tag using the Gibson assembly master mix (New England Biolabs) for *Ta*SS4-1B, or BamHI and XhoI sites for *Ta*BGC1-4B. Proteins were expressed in *E. coli* strain BL21 as described in Seung *et al*. (2015). Denaturing purification of the protein with urea was carried out using the Ni-NTA Agarose (Qiagen). Immunisation of rabbits was carried out at Eurogentec. Antibodies were enriched from antiserum using protein A-agarose (Sigma-Aldrich). Affinity purification of *Ta*BGC1-specific antibodies from the antiserum was performed with a HiTrap NHS-Activated HP column (GE Healthcare), conjugated to *Ta*BGC1 recombinant protein.

For immunoblotting: endosperms from developing grains were dissected and homogenised in 40 mM Tris-HCl, pH 6.8, 5 mM MgCl_2_, 2% *(w/v)* SDS, protease inhibitor cocktail (Roche). The homogenate was heated at 95°C for 10 min, and insoluble material was removed by centrifugation at 20,000*g* for 10 min. The concentration of proteins was determined using the BCA assay (Thermo Scientific). The following dilutions of primary antibodies were used for immunoblotting: anti-*Ta*SS4: 1:200, anti-*Ta*BGC1: 1:200, anti-actin (Sigma-Aldrich; A0480): 1:10,000, anti-YFP (Torrey pines; TP401): 1:5,000, or anti-HA (Abcam; ab9110): 1:5,000. Bands were detected using the IRDye 800CW-donkey-anti-rabbit or 680RD-donkey-anti-mouse secondary antibodies (1:10,000; Li-Cor) and the Odyssey Classic Imaging system (Li-Cor).

## RESULTS

### Mutants lacking both TaSS4 homoeologs produce aberrant endosperm starch

Hexaploid wheat has three homoeologs of *TaSS4* on group 1 chromosomes (Irshad *et al*., 2019). The B- and D-genome homoeologs were reported to have 16 exons and the A-genome homoeolog has 13 exons. We established that in the most recent wheat genome release (RefSeq v1.1 cv. Chinese Spring; Appels *et al*., (2018)), these homoeologs correspond to *TaSS4-1A* (TraesCS1A02G353300), *TaSS4-1B* (TraesCS1B02G368500) and *TaSS4-1D* (TraesCS1D02G356900)(Fig. 1a). As reported by Irshad *et al*. (2019), *TaSS4-1B* and *TaSS4-1D* loci contained 16 exons (Fig. 1b) but *TaSS4-1A* had a shorter coding sequence generated from 13 exons. Nonetheless, the predicted transcript length of *TaSS4-1A* was the same as that of the other homoeologs because it had a longer 5′ UTR. To investigate the discrepancy in gene model between homoeologs, we compared nucleotide and predicted amino acid sequences from the Chinese Spring reference sequence with those of other sequenced hexaploid wheat cultivars (Cadenza, Paragon, Robigus, Claire) and the tetraploid cultivar Kronos on the Grassroots database (Clavijo *et al*., 2017)(Fig. S1a). For all cultivars except for Chinese Spring, *TaSS4-1A* was predicted to have all 16 coding exons. Chinese Spring had a unique single nucleotide polymorphism (SNP) that results in a premature stop codon in a position occupied by exon 4 in the gene model for the other cultivars (Fig. 1b,S1a). This SNP most likely led to the incorrect prediction of 13 coding exons and a long 5′ UTR for *TaSS4-1A* in the Chinese Spring sequence.

To assess the importance of *Ta*SS4 in endosperm starch formation, we created mutants of tetraploid wheat that are defective in both homoeologs of *Ta*SS4. We used the TILLING collection of exome-capture sequenced, EMS-mutagenized lines of the tetraploid wheat Kronos (Krasileva *et al*., 2017; http://www.wheat-tilling.com) to identify mutants that are likely to have no *Ta*SS4-1A or *Ta*SS4-1B protein. The predicted amino acid sequences of *TaSS4-1A* and *TaSS4-1B* from Kronos shared 99-100% identity with those from the Chinese Spring reference genome, and the two Kronos homoeologs were 97% identical to each other (Fig. S1a,b). For *TaSS4-1A*, we obtained the Kronos2166(K2166) line, which carries a splice donor site mutation after exon 5 (Fig. 1b). For *TaSS4-1B*, we obtained Kronos2565(K2565) carrying a premature stop codon in place of Trp364, and Kronos1450(K1450) carrying a splice acceptor site mutation before exon 15. The presence of each mutation was confirmed by KASP genotyping, using the primers in Table S1. The K2166 line was crossed with K2565 to create the *Tass4-1* lines, or with K1450 to create the *Tass4-2* lines. KASP genotyping was used to identify F_2_ individuals homozygous for both A and B mutations (*aa bb*), homozygous for only the *TaSS4-1A* mutation (*aa* BB) or the *TaSS4-1B* mutation (AA *bb*), and ‘negative segregant’ controls that lacked both mutations (AA BB). Except where specified, observations below were on the *Tass4-1* lines.

To observe *Ta*SS4 protein levels during grain development and the effect of the *Tass4-1* mutations on *Ta*SS4 protein abundance, we generated an antiserum against a *Ta*SS4-1B recombinant protein, expressed in and purified from *Escherichia coli*. Immunoblots of *Ta*SS4-1A and *Ta*SS4-1B proteins transiently expressed in *Nicotiana benthamiana* leaves demonstrated that the antiserum recognised both homoeologs (Fig. S2). Protein extracts from endosperms dissected from wild-type developing grains (10, 15 and 20 days post anthesis (dpa)) were immunoblotted with the antiserum. A band that corresponded to the predicted size of the mature polypeptide (98 kDa) was observed at all three timepoints, but was most prominent at the 10 dpa timepoint (Fig. 2a). Several other bands at different molecular weights were detected, but comparison of immunoblots from the wild-type and mutant extracts showed that the 98 kDa band was missing in the latter while other bands were unaffected (Fig. 2b). We conclude that the 98 kDa band represents *Ta*SS4, and that the other bands result from non-specific binding.

**Fig. 2.**
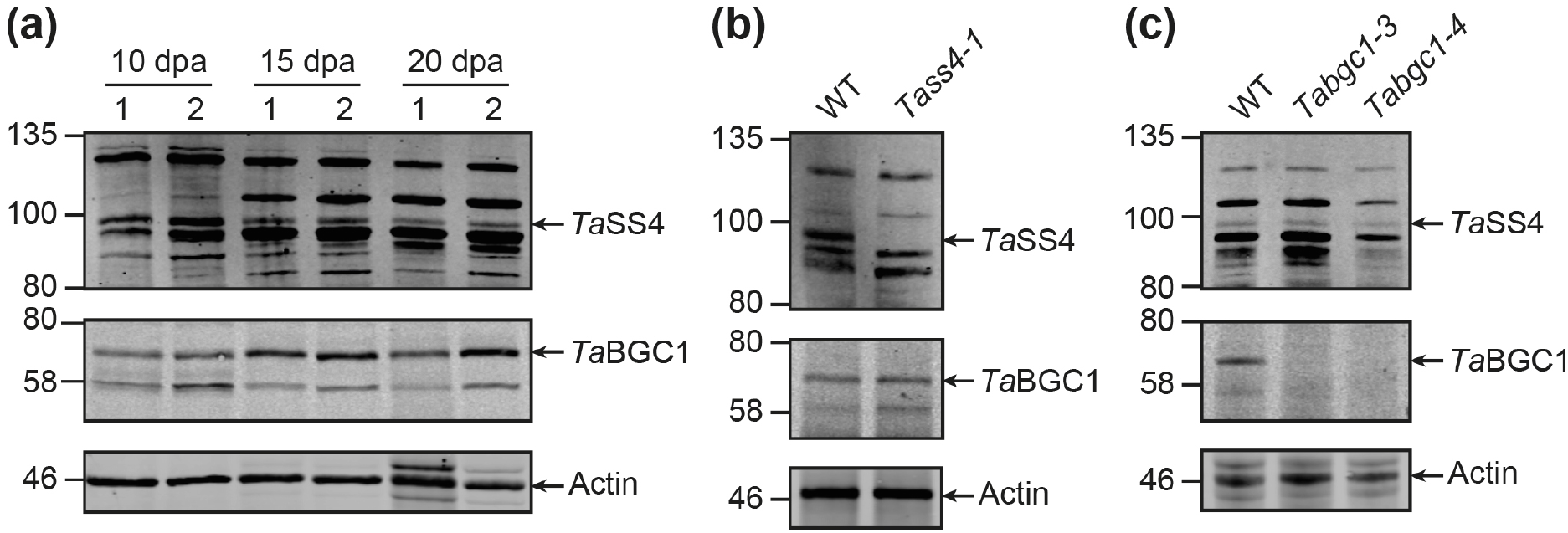
*Ta*SS4 and *Ta*BGC1 protein levels in developing endosperm. **(a)** Total proteins were extracted from developing endosperms at 10, 15 and 20 dpa, and immunoblotted using anti-*Ta*SS4 (upper panel), anti-*Ta*BGC1 (middle panel) or anti-actin (lower panel) antibodies. Lanes were loaded on an equal protein basis. The migration of molecular weight markers are indicated in kilodaltons (kDa) to the left of each panel. Two replicate extractions for each genotype (numbered 1 and 2) were prepared from grains harvested from two different plants. **(b)** Same as **(a)**, but with *Tass4-1* grains harvested at 10 dpa. **(c)** Same as **(a)**, but with *Tabgc1* grains harvested at 20 dpa.

Transcript data for whole caryopses (Maccaferri *et al*., 2019) and dissected endosperm (Xiang *et al*., 2019) from developing tetraploid wheat grains revealed that *TaSS4* transcript levels were higher at the early stages of grain development (8-11 dpa) than later stages (16-22 dpa)(Fig. S3). These data are consistent with the observed decrease in *Ta*SS4 protein levels at later stages of grain development (Fig. 2a).

To assess the impact of the *Tass4-1* mutations on endosperm starch, we purified starch granules from mature grains and observed them using scanning electron microscopy. Granules from control lines (AA BB) and the single homoeolog mutants (*aa* BB and AA *bb*) had flattened A-type granules and round B-type granules typical of wheat starch (Fig. 3a). By contrast, most starch granules from the double mutant (*aa bb*) had irregular, polyhedral morphology. The irregular granules were highly variable in size, but rarely exceeded the size of a typical A-type granule. A-type granules of normal appearance were also present in the double mutant, but we rarely observed normal B-type granules.

**Fig. 3.**
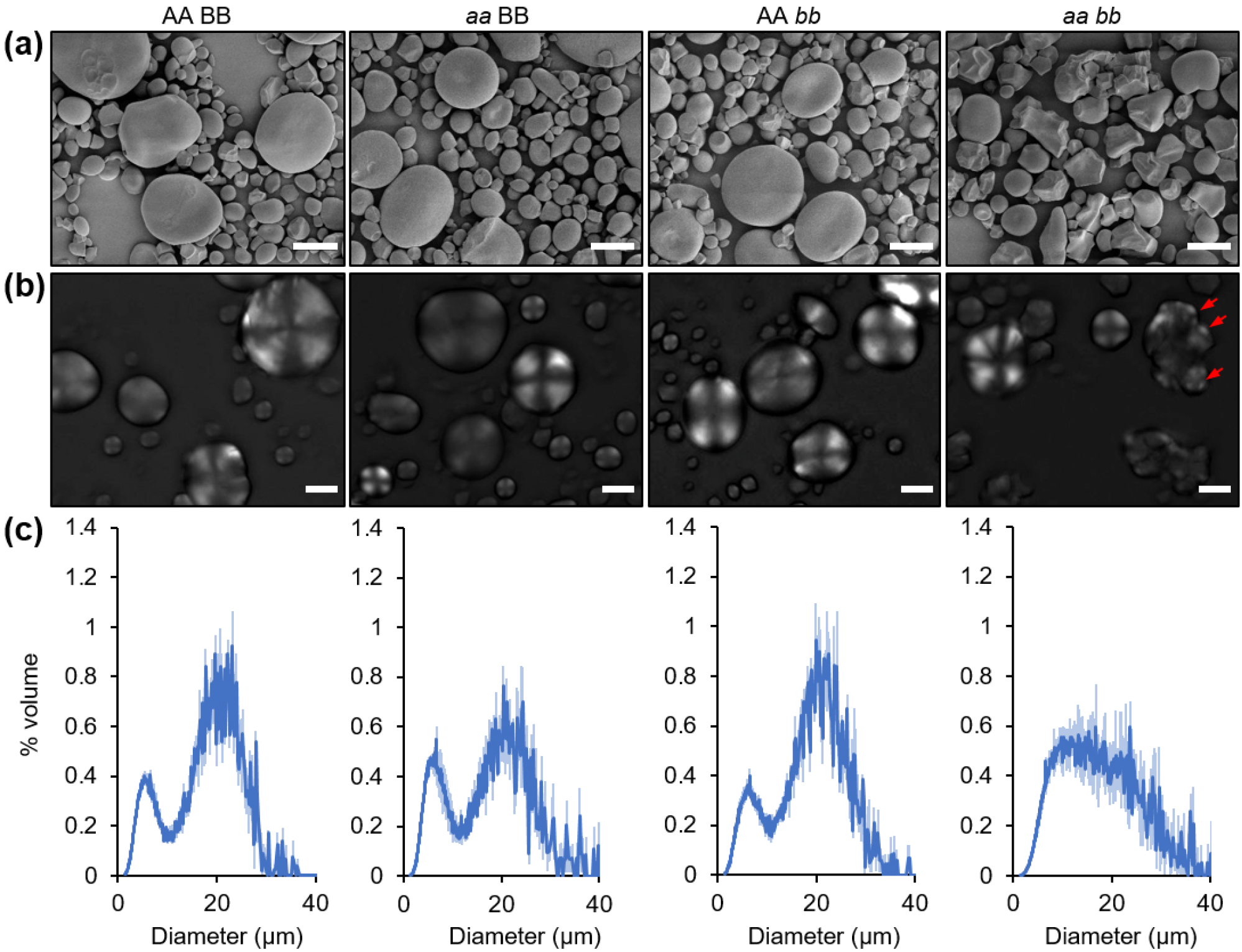
*Tass4-1* double mutants have severely altered granule morphology. **(a)** Endosperm starch granules from mature grains were observed using scanning electron microscopy. Single (*aa* BB or AA *bb*) and double mutants (*aa bb*) were compared with control (AA BB) lines. Bars = 10 μm. **(b)** As **(a)**, but granules were observed using cross-polarised light microscopy. The multiple hila in the large polyhedral granule are indicated with red arrows. Bars = 10 μm. **(c)** Size distribution of endosperm starch granules. The volume of granules at each diameter relative to the total granule volume was quantified using a Coulter counter. Values represent mean (solid line) ± SEM (shading) of three replicate starch extractions from grains of three different plants.

We used cross-polarised light microscopy to examine the origins of the larger polyhedral granules in the *Tass4-1* double mutant endosperm. In the control line and the single homoeolog mutants, there was one ‘Maltese cross’ per A-type or B-type granule, indicating a single centre of organisation (Fig. 3b). The few normal A-type granules in the double mutant also had single crosses. However, a complex birefringence pattern with faint or multiple crosses were observed in most of the polyhedral granules, indicating multiple initiation points.

Using a Coulter counter, we examined the granule size distribution in the endosperm starches. As expected, starch from the control line and single mutants showed a bimodal size distribution, with peaks at approx. 20 μm diameter for A-type and 7-8 μm diameter for B-type granules (Fig. 3c). The size and relative proportion (by volume) of A-type and B-type granules were quantified by fitting a mixed log-normal distribution (Table 1; Tanaka *et al*., 2017). There were no significant differences between the control and single mutants. The granule size distribution of the double mutant had no distinct peaks, and neither a mixed nor a single distribution could be fitted reliably to these data.

**Table 1:**
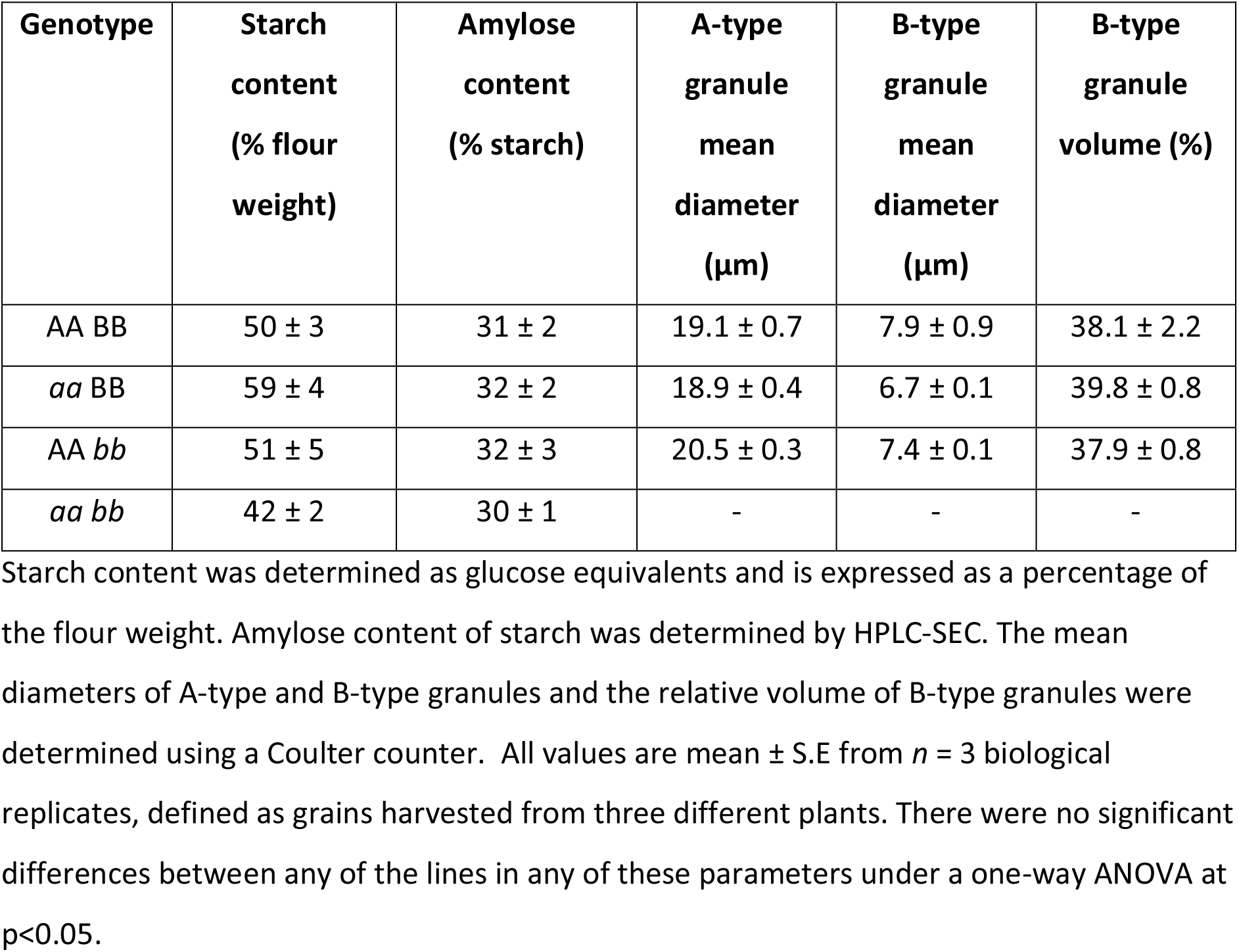
Starch content, composition and granule size in *Tass4-1* mature grains.

| Genotype | Starch content (% flour weight) | Amylose content (% starch) | A-type granule mean diameter (μm) | B-type granule mean diameter (μm) | B-type granule volume (%) |
| --- | --- | --- | --- | --- | --- |
| AA BB | 50 ± 3 | 31 ± 2 | 19.1 ± 0.7 | 7.9 ± 0.9 | 38.1 ± 2.2 |
| aa BB | 59 ± 4 | 32 ± 2 | 18.9 ± 0.4 | 6.7 ± 0.1 | 39.8 ± 0.8 |
| AA bb | 51 ± 5 | 32 ± 3 | 20.5 ± 0.3 | 7.4 ± 0.1 | 37.9 ± 0.8 |
| aa bb | 42 ± 2 | 30 ± 1 | - | - | - |

Starch content was determined as glucose equivalents and is expressed as a percentage of the flour weight. Amylose content of starch was determined by HPLC-SEC. The mean diameters of A-type and B-type granules and the relative volume of B-type granules were determined using a Coulter counter. All values are mean ± S.E from $n = 3$ biological replicates, defined as grains harvested from three different plants. There were no significant differences between any of the lines in any of these parameters under a one-way ANOVA at $p < 0.05$ .

The normal granule morphology of control (AA BB) and single mutant (*aa* BB and AA *bb*) lines indicates that the aberrant morphology arises only when both *Tass4* homoeologs are defective (*aa bb*). However, it remained possible that the aberrant morphology arose from a combination of background mutations in the single-mutant parents of the double mutant. To exclude this possibility, we first backcrossed the double mutant to the wild type twice, and re-isolated the double mutant in the BC_2_F_2_ generation. Aberrant granule morphology was still observed after the backcrosses (Fig. S4). Granule size distributions of backcrossed and non-backcrossed *Tass4-1 aa bb* lines were identical, indicating that this phenotype is unlikely to arise from background mutations. Second, we examined granule morphology in a second set of mutant lines, *Tass4-2*, obtained by crossing K2166 with an independent mutant for *TaSS4-1B*, K1450 (see above, Fig. 1b). The *Tass4-2 aa bb* double mutant had the same aberrant granule morphology as the *Tass4-1* lines (Fig. S4).

### TaSS4 mutations do not alter total starch content, composition or amylopectin structure

We investigated whether the aberrant granule morphology in the *Tass4-1 aa bb* line was accompanied by changes in starch content, composition or structure. The starch content of mature grains was not significantly different on a dry weight basis between control, single and double mutant lines (Table 1). To examine amylopectin/amylose structure and abundance, debranched starch was subjected to High Performance Liquid Chromatography-Size Exclusion Chromatography (HPLC-SEC) with refractive index detection (Cave *et al*., 2009; Tuncel *et al*., 2019). The chain length distribution of amylopectin and amylose, and the estimated amylose content, were identical in control and mutant starches (Fig. S5, Table 1). Thus, the altered granule morphology in the *Tass4-1* double mutant cannot be attributed to differences in starch content, composition, or polymer structure.

### The Tass4 mutant shows defective granule morphology during early grain development

To investigate at which stage of grain development the aberrant granules in the *Tass4-1* double mutant form, we examined granule morphology and size distribution of starch extracted from developing endosperms at three different time points: 8, 14 and 21 dpa. As expected, the wild-type endosperm contained only A-type granules at 8 dpa, with a peak at approx. 15 μm diameter (Fig. 4a,b). B-type granules were present at 14 and 21 dpa, with a peak at approx. 5 μm diameter. Starch from the *Tass4-1* double mutant already contained aberrant, polyhedral granules at 8 dpa, and no distinct A-type and B-type granule peaks were observed in the mutant at any timepoint.

**Fig. 4.**
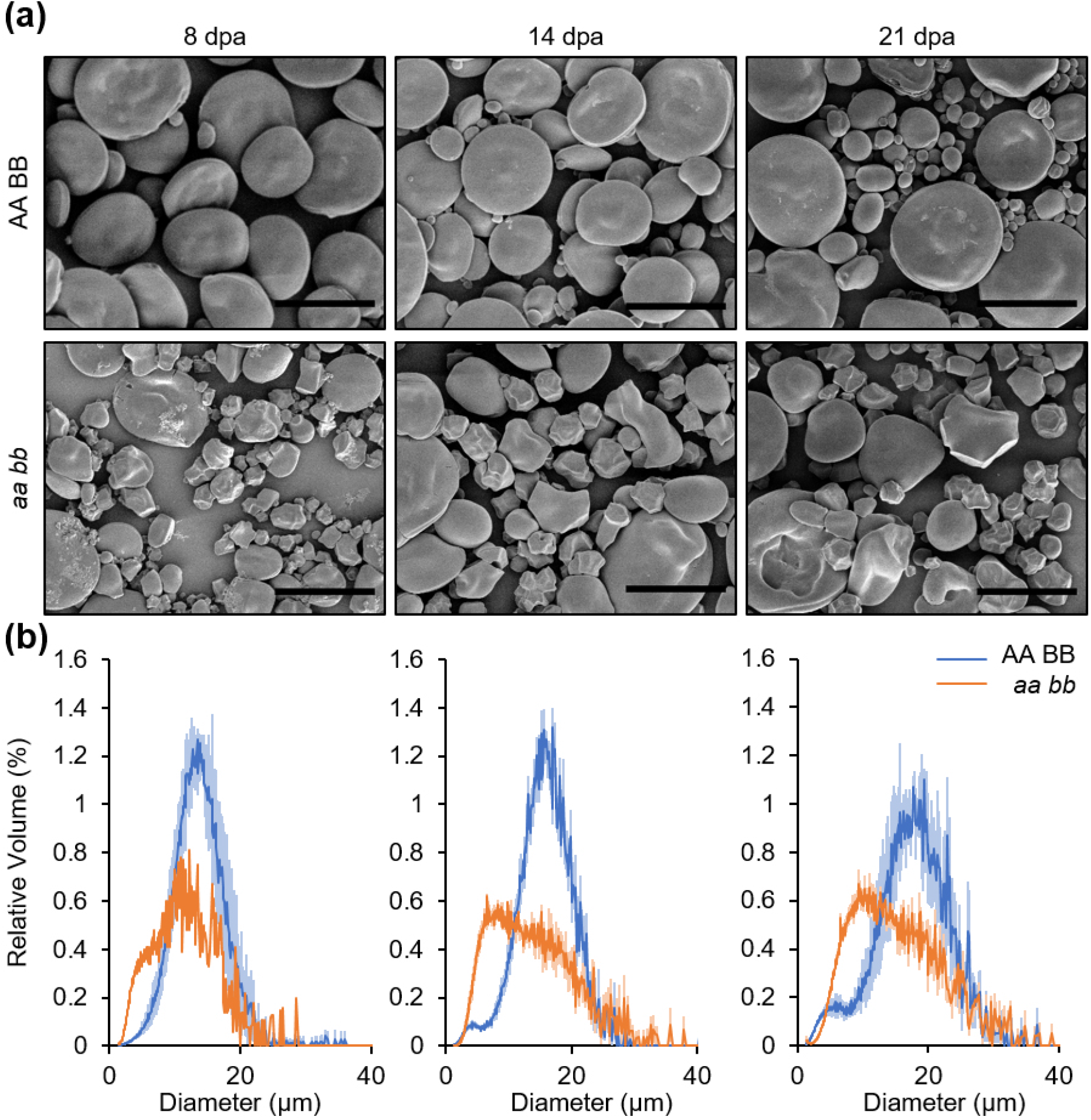
Aberrant granules in *Tass4-1* double mutants form early in grain development. **(a)** Endosperm starch from developing grains (8, 14 and 21 dpa) of the *Tass4-1* double mutant (*aa bb*) and control lines (AA BB) were observed using scanning electron microscopy. Bars = 25 μm. **(b)** Size distribution of endosperm starch granules. The volume of granules at each diameter relative to the total granule volume was quantified using a Coulter counter. Values represent mean (solid line) ± SEM (shading) of three replicate starch extractions from grains of three different plants, except for *aa bb* at 8 dpa where one starch extraction was performed.

### The Tass4 mutant produces compound granules

We examined the spatial arrangement of the polyhedral granules within the endosperm of the *Tass4-1* double mutant, initially in thin sections of mature grains stained with iodine solution. Consistent with the observations made on purified starch, the control and single mutant lines had normal A- and B-type granules. The granules with polyhedral shapes in the double mutant were almost always tessellated within larger structures (Fig. 5).

**Fig. 5.**
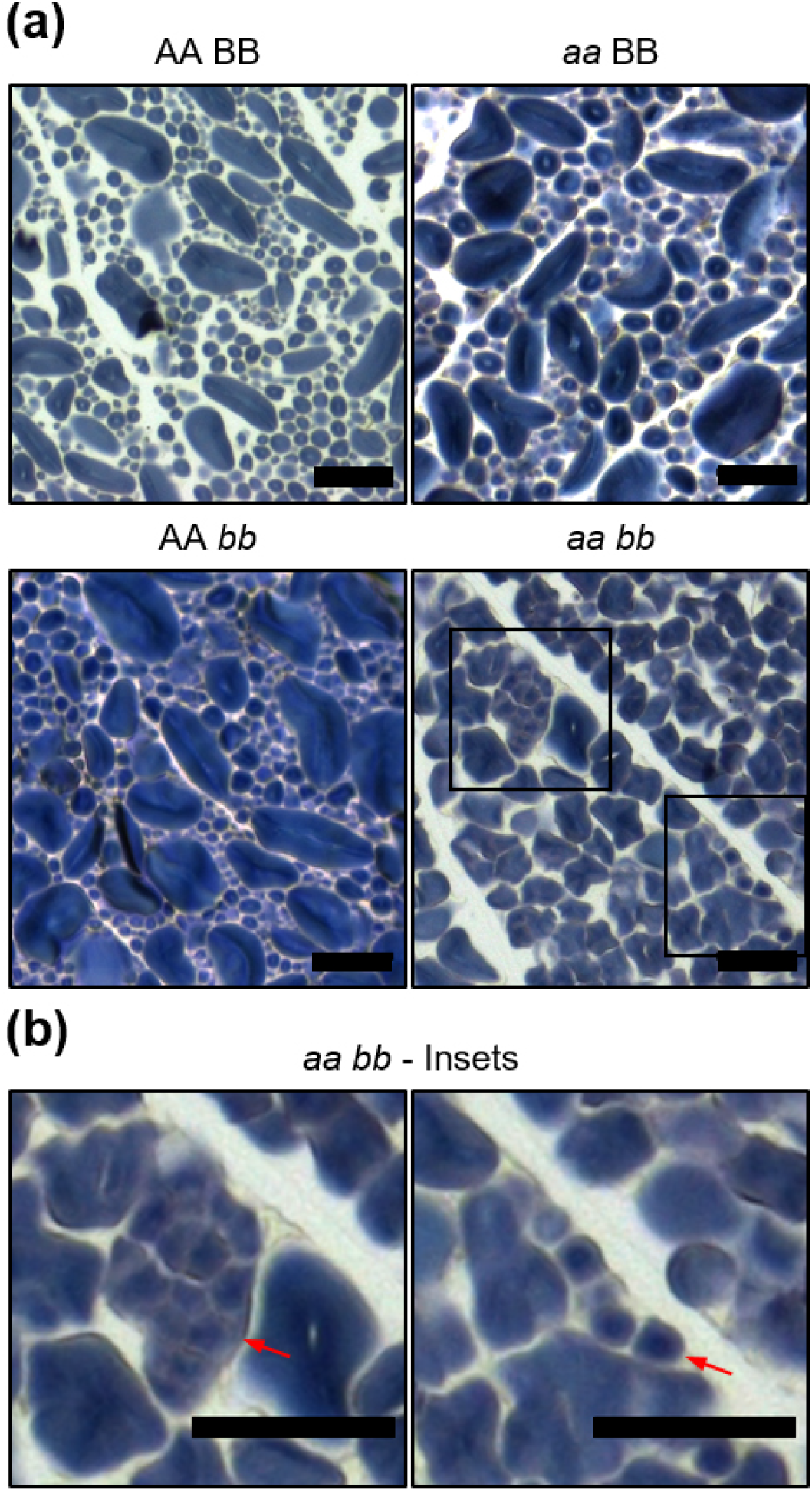
Endosperm sections of *Tass4-1* single and double mutants. **(a)** Thin sections were prepared from mature grains, stained with iodine and observed using light microscopy. Single (*aa* BB or AA *bb*) and double mutants (*aa bb*) were compared with control (AA BB) lines. Bars = 25 μm. **(b)** Insets showing a close-up view of a large compound structure (left panel) and a tubule-like structure (right panel) in the *aa bb* section – both indicated with red arrows. Bars = 25 μm.

Observing sections of developing endosperm at 15 dpa by light microscopy and TEM revealed remarkable heterogeneity among amyloplasts in the mutant (Fig. 6). Whereas the amyloplasts in the wild type contained single A-type granules and some peripheral stroma, most amyloplasts in the mutant contained compound granules, while some contained A-type granules that were indistinguishable from those of the wild type - and in most endosperm cells, both types of amyloplasts were present. The number of individual ‘granulae’ visible within each compound granule section varied, ranging from 4 to >40. Notably, some amyloplasts had formed granules that were tessellated in tubular structures.

**Fig. 6.**
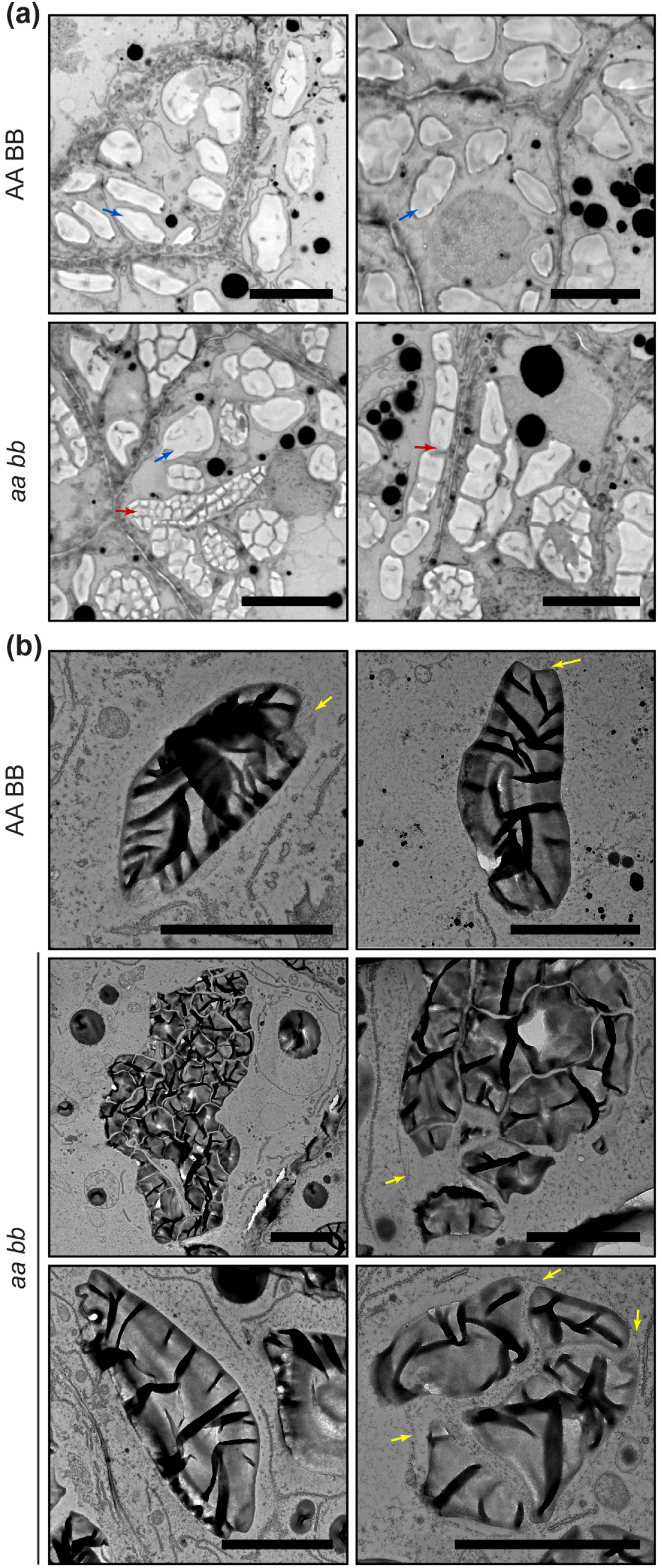
Compound granules in the developing *Tass4-1* endosperm. **(a)** Toluidine blue-stained sections of developing endosperm (15 dpa) in the *Tass4-1* double mutant (*aa bb*) or control (AA BB), observed using light microscopy. Blue arrows indicate examples of normal A-type granules, while red arrows indicate compound granules. Bars = 20 μm. **(b)** Same as **(a)**, but observed using transmission electron microscopy. Amyloplast membranes and stromal space around granules are indicated with yellow arrows. Bars = 5 μm.

### Tass4 starch granules resemble those of the Tabgc1 mutant

The elimination of another component of the putative granule initiation complex defined in Arabidopsis, PTST2/FLO6/BGC1, results in strong granule morphology defects in endosperms of barley and hexaploid wheat, including the occurrence of “semi-compound” granules (Suh *et al*., 2004; Chia *et al*., 2020). To discover the relationship between the roles of *Ta*SS4 and *Ta*BGC1 in wheat endosperm, we compared phenotypes of *Tass4* and *Tabgc1* mutants in the same Kronos background, and tested whether *Ta*SS4 and *Ta*BGC1 proteins can interact with each other. For two *Tabgc1 aa bb* double mutants (Fig. 7a) that accumulate no detectable *Ta*BGC1 protein (Fig. 2c), we established that starch granules had morphologies like those described for *Tabgc1* mutants in hexaploid wheat and barley: mature grains contained A-type granules of normal appearance and small polyhedral granules (Fig. 7b). As in the *Tass4* mutant, these polyhedral granules were already present during early grain development (8 dpa onwards)(Fig. S6). However, normal A-type granules were more frequent in the *Tabgc1* mutant (Fig. 7b) than in the *Tass4* mutant (Fig. 3a). Coulter counter analysis also showed a prominent A-type granule peak in the *Tabgc1* mutant at a similar diameter to that of the wild type (Fig. 7c, d). Such a distinct peak was not observed in the *Tass4-1* mutant (Fig. 3c).

**Fig. 7.**
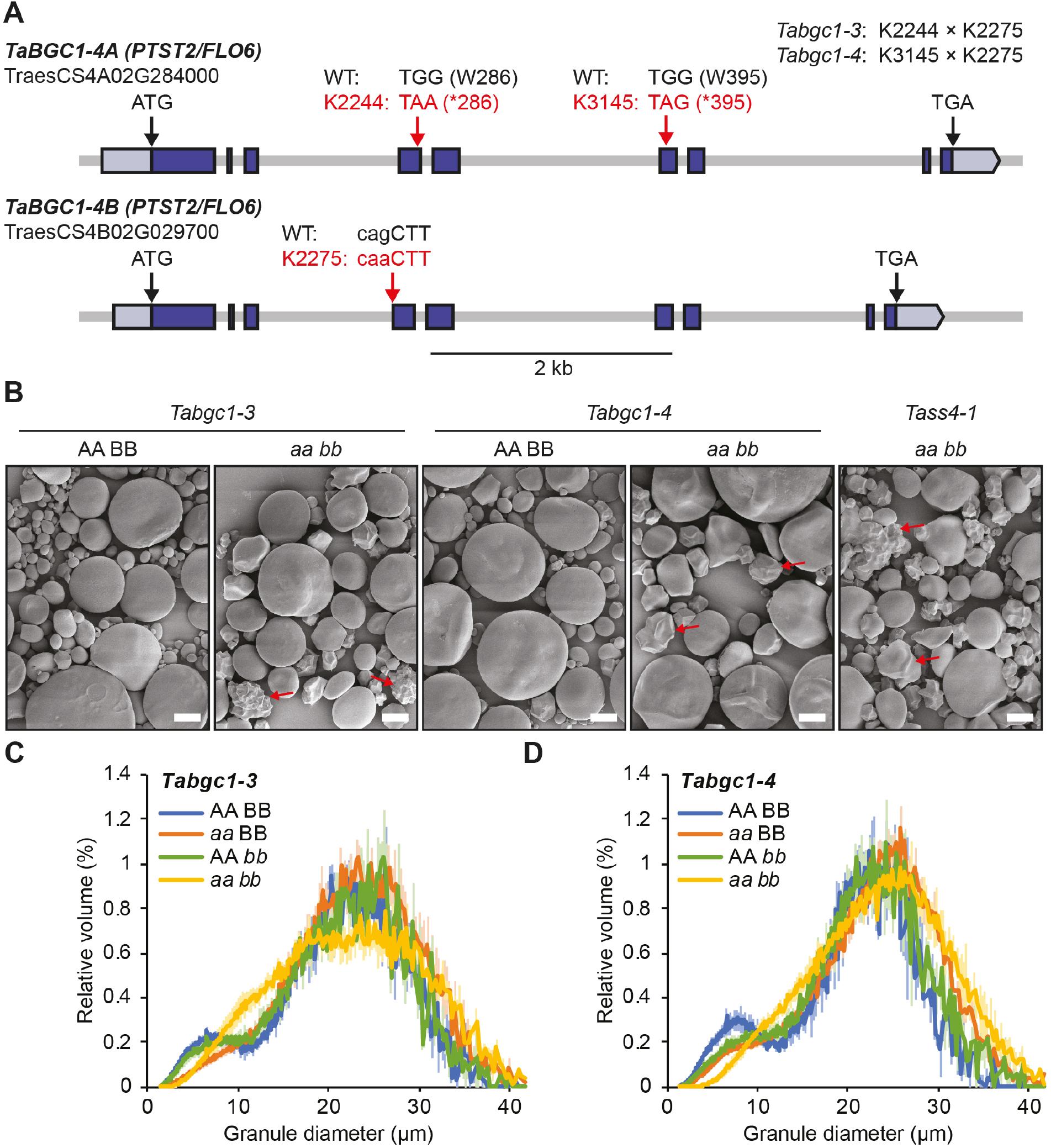
Similar defects in granule morphology in *Tass4* and *Tabgc1* mutants. **(a)** Gene models of the *TaBGC1* homoeologs. Exons are represented with blue boxes, while light blue boxes represent the 5′ and 3′ UTRs. The locations of the mutations in the *Tabgc1-3* and *Tabgc1-4* lines are depicted with red arrows and the mutated codons/amino acids are shown in red letters. **(b)** Purified starch granules from mature grains of the double mutants (*aa bb*) and control lines (AA BB) observed using scanning electron microscopy. Examples of polyhedral granules are marked with red arrows. Bars = 10 μm. **(c)** Size distribution of endosperm starch granules in mature grains of the *Tabgc1-3* single (*aa* BB and AA *bb*) and double mutants. The volume of granules at each diameter relative to the total granule volume was quantified using a Coulter counter. Values represent mean (solid line) ± SEM (shading) of three replicate starch extractions from grains of three different plants. **(d)** Same as **(c)**, but with *Tabgc1-4*.

### Loss of TaSS4 affects starch synthesis in pollen

The *Tass4* double mutant was indistinguishable from control lines in terms of plant growth (Fig. 8a). Most grains of the mutant appeared normal and the average weight of individual grains was not significantly altered compared to the wild type, although we observed rare examples of smaller, shrivelled grains in the mutant (Fig. 8b, S7). While the double mutant produced comparable numbers of tillers to the control (Fig. 8c), the number of grains per spike was significantly reduced in the mutant (Fig. 8d). This reduction in grain number was most severe in the non-backcrossed *Tass4-1* double mutant but was partly recovered after backcrossing, suggesting that this phenotype was exacerbated by background mutations in non-backcrossed lines (Fig. 8d). Since the fewer grains in the backcrossed mutant suggests defective fertilisation, we examined starch accumulation in pollen grains of the mutant using iodine staining. Less than a third of pollen grains from the double mutant contained starch, contrasting those from control lines where almost all contained starch (Fig. 8e, S8a). Cross-pollination experiments with the backcrossed *Tass4-1* lines demonstrated that using *aa bb* pollen to fertilise AA BB maternal plants resulted in significantly reduced fertilisation rates compared to the reciprocal cross (Fig. S8b). These data suggest that *Ta*SS4 is important for normal pollen starch synthesis and viability. Interestingly, grains from low-yielding non-backcrossed and high-yielding backcrossed *Tass4-1* lines had identical starch granule morphology (Fig. S3), demonstrating that this phenotype is independent from the fertility phenotype.

**Fig. 8.**
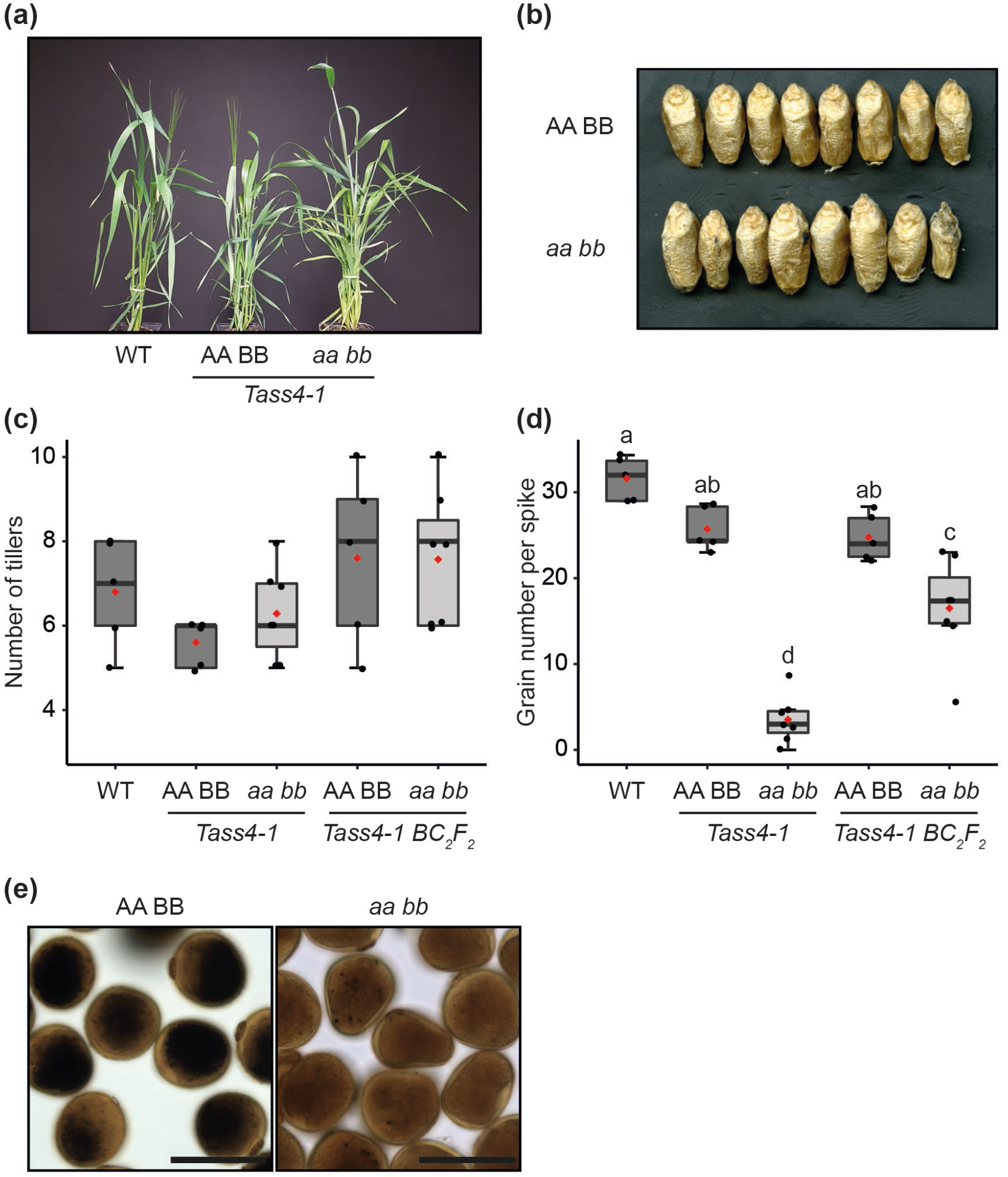
Growth and fertility phenotypes of the *Tass4* mutant. **(a)** Photograph of 8-week old plants of wild type, *Tass4-1* control (AA BB) and double mutant (*aa bb*). **(b)** Photographs of mature grains. **(c)** The number of tillers on backcrossed (BC_2_F_2_) and non-backcrossed *Tass4-1* mutants were counted on *n*=5-7 plants. Individual data points (black dots) and the mean (red dot) are shown over the box plots. There were no significant differences between the lines under a one-way ANOVA. **(d)** The average number of grains in the three primary spikes was calculated for n=5-7 plants. Plots are as for **(c)**. Different letters indicate significant differences at p < 0.05 under a one-way ANOVA and Tukey’s posthoc test. Note that the statistical analysis includes data in Fig. S8c, which were obtained in the same experiment. **(e)** Iodine-stained pollen grains observed with light microscopy. Bars = 50 μm.

### Loss of TaSS4 results in fewer starch granules per leaf chloroplast

Since SS4 plays a critical role in granule initiation and morphogenesis in Arabidopsis leaves, we investigated whether these roles are conserved in wheat leaves. Leaves of the *Tass4-1* double mutant accumulated less than half the starch content of the control over the light period (Fig. 9a). Light microscopy to visualise granules in chloroplasts at the end of the day showed similar frequency distributions of granule sections per chloroplast section in the control and single mutant lines: 70 to 80% of chloroplasts contained between 1-8 granule sections and the remainder contained no visible starch granule (Fig. 9b, c). By contrast, almost 80% of chloroplasts in the double mutant contained no visible starch granule. Examination with TEM showed that granules in control leaves had the typical flattened shape of leaf starch, whereas most granules in the double mutant were small and rounded (Fig. 9d).

**Fig. 9.**
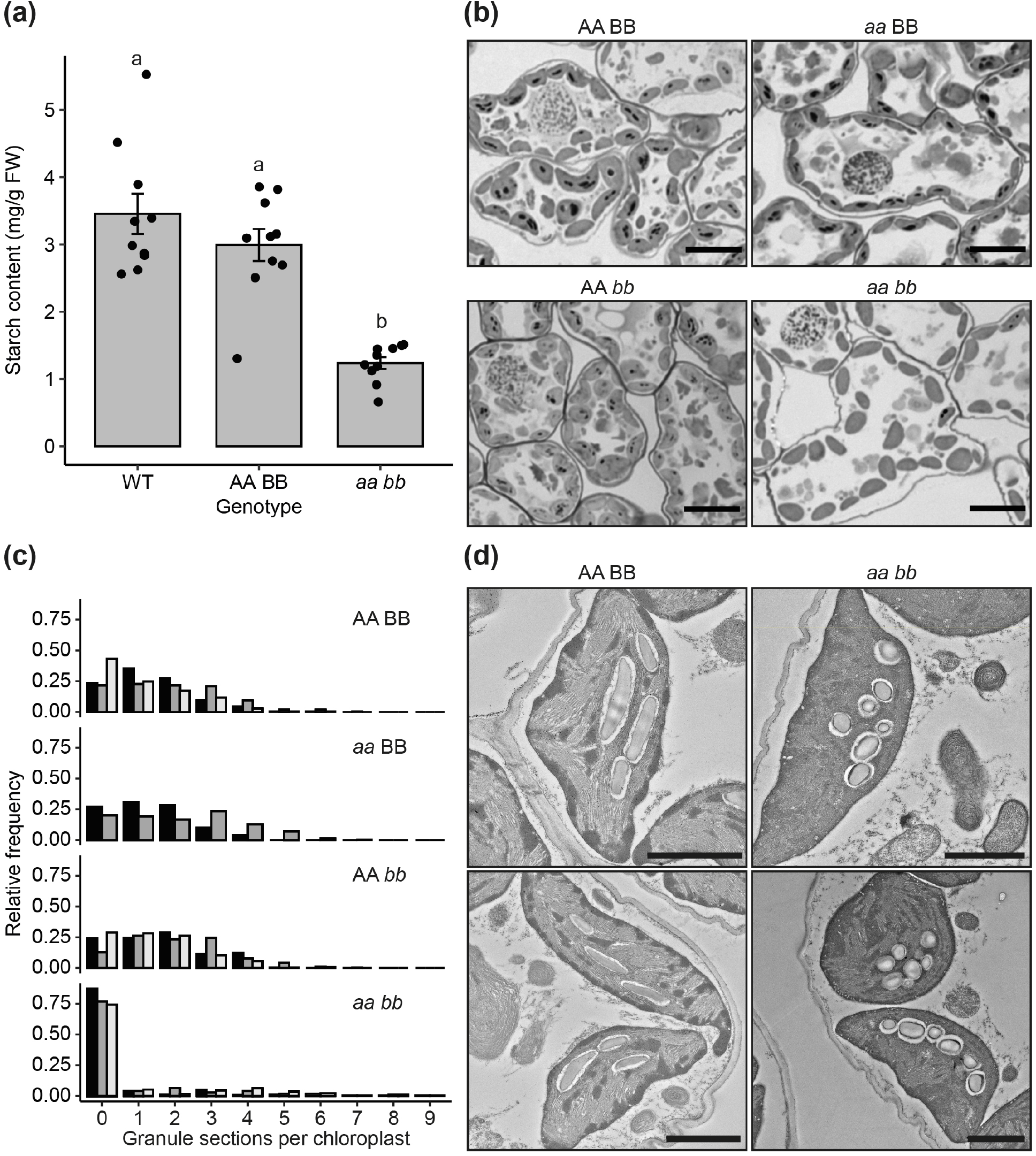
Loss of *TaSS4* results in fewer starch granules in leaf chloroplasts. **(a)** Leaf starch content at the end of day. Bars represent the mean ± SEM from *n* = 10 plants. Different letters indicate significant differences at p<0.05 under a one-way ANOVA and Tukey’s posthoc test. **(b)** Starch granules in mesophyll chloroplasts observed with light microscopy. Leaf samples were harvested at the end of day from the middle of the flag leaf of 5-week old plants. Thin sections were stained with toluidine blue and periodic acid/Schiff’s reagent. Bars = 10 μm. **(c)** Quantification of starch granule number per chloroplast. Three replicate sections for each genotype (each produced from separate plants, plotted as black, dark grey and light grey bars) were observed using light microscopy as in **(b)** (except for *aa* BB, where two replicate sections were produced). Histograms represent the frequency of chloroplast sections containing a given number of granule sections, relative to the total number of chloroplast sections. A total of 217-237 chloroplasts were analysed for each replicate. **(d)** Leaf chloroplasts were imaged using transmission electron microscopy. Bars = 2 μm.

These results suggest that as in Arabidopsis, the loss of *Ta*SS4 in wheat strongly affects the number of granules initiated per chloroplast. We therefore attempted to complement the Arabidopsis *Atss4* mutant by expression of *Ta*SS4-1B with a C-terminal YFP tag and under the Arabidopsis Ubiquitin 10 promoter (pUBQ). The *Atss4-1* mutant had pale leaves, but the transformed lines were not pale (Fig. 10a). The transformed lines had multiple granules in most chloroplasts, whereas most chloroplasts of the *Atss4* mutant were either starchless or contained a single large, round granule (Fig. 10b). *Ta*SS4 can thus partially complement the granule number phenotype of the *Atss4* mutant. Most granules in the transgenic lines were also irregularly shaped, and few were flattened as in the wild-type, or round as in *Atss4* (Fig. 10c) - suggesting *Ta*SS4 can also influence granule morphology when expressed in Arabidopsis leaves.

**Fig. 10.**
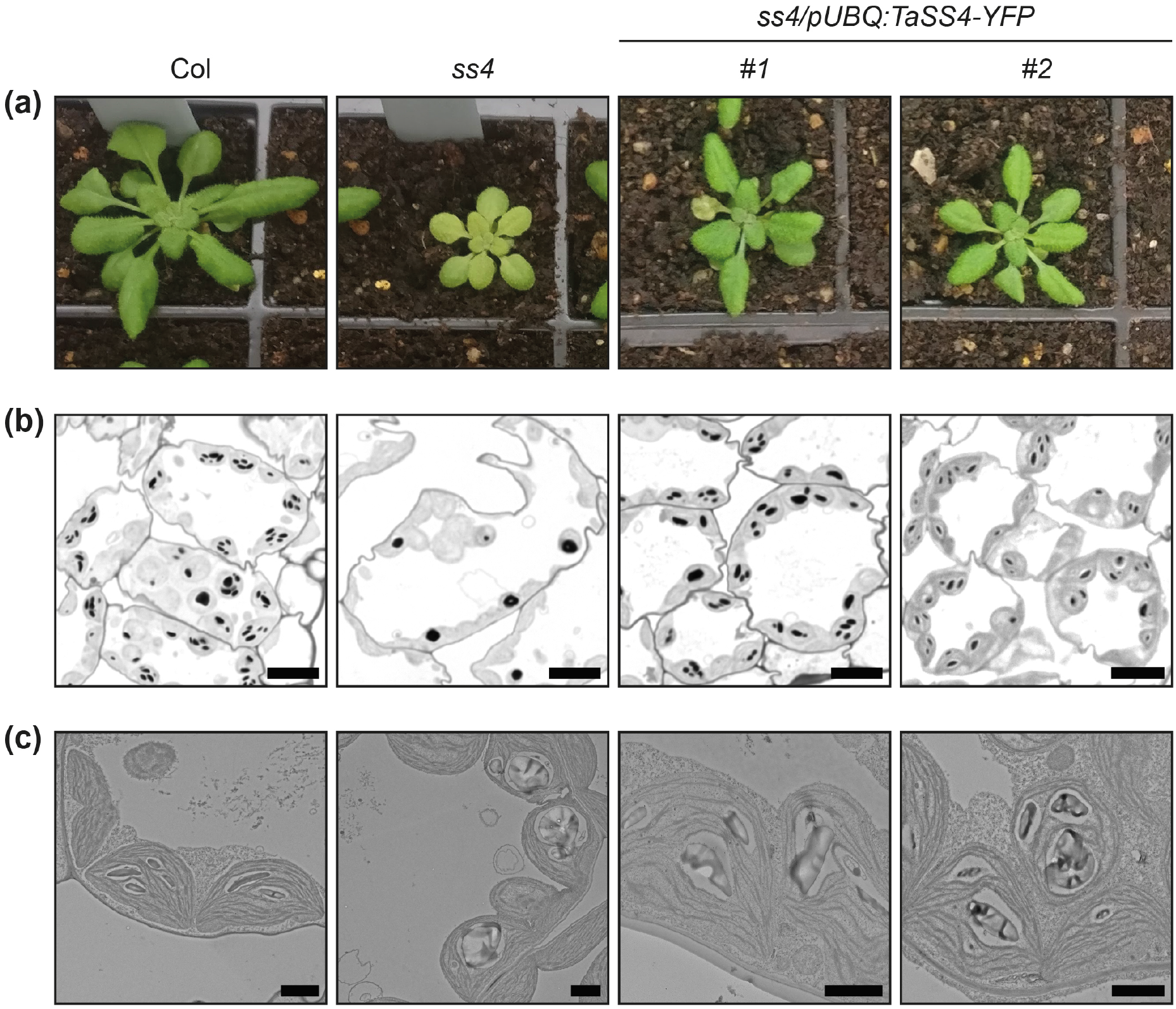
*Ta*SS4 can partially complement plant growth and starch granule morphology phenotypes of the Arabidopsis *Atss4* mutant. **(a)** Rosette morphology of the wild type (Col-0), *Atss4* mutant, and two independent transgenic lines expressing *Ta*SS4 1B-YFP under the Arabidopsis *Ubiquitin10* promoter in the *Atss4* mutant background (*ss4/pUBQ:TaSS4-YFP*). **(b)** Starch granules in chloroplasts observed using light microscopy for the plants shown in **(a)**. Thin sections of young leaves of a 5-to 6-week old rosette were stained using toluidine blue and periodic acid/Schiff’s reagent. Bars = 10 μm. **(c)** Same as **(b)**, but viewed under transmission electron microscopy. Bars = 2 μm.

## DISCUSSION

### TaSS4 is necessary for normal granule initiation in the endosperm

In wheat endosperm, granule initiation is spatially and temporally coordinated such that single A-type granules form in amyloplasts during early grain development and B-type granules initiate later and at least partially in stroma-filled tubules (stromules) that emanate from the amyloplast (Parker, 1985; Langeveld *et al*., 2000). This pattern is distinct from most other grasses (e.g: rice), which form compound granules by initiating multiple granules per amyloplast during early grain development (Matsushima *et al*., 2013, 2015). Recent work in Arabidopsis leaves has suggested a mechanism of granule initiation in leaf chloroplasts involving at least six proteins – SS4, SS5, PTST2, PTST3, MFP1 and MRC, each of which is individually necessary for normal granule initiation (Seung & Smith, 2019; Abt & Zeeman, 2020). Among these initiation proteins, only SS4 is known to have enzymatic activity (Roldán *et al*., 2007; Szydlowski *et al*., 2009; Abt *et al*., 2020). However, the influence of SS4 on the distinct granule initiation patterns observed in cereal amyloplasts was not known. Our study demonstrates that *Ta*SS4 is required for the control of granule initiation in wheat endosperm. Loss of *Ta*SS4 in wheat did not affect the content, composition or polymer structure of endosperm starch (Table 1), but resulted in the formation of compound granules in the endosperm in place of most A-type granules (Fig. 3–6). A similar phenotype was observed in mutants fully deficient in *Ta*BGC1 in tetraploid wheat (Fig. 7), and in hexaploid wheat (Chia *et al*., 2020), suggesting that the two proteins act in a similar process. However, the *Tass4* phenotype was more severe than the *Tabgc1* phenotype as there were substantially more normal A-type granules in the latter (Fig. 7). These observations parallel those in Arabidopsis leaves, in which granule initiation is more compromised in the *Atss4* mutant than in the *Atptst2* mutant (Seung *et al*., 2017).

To our knowledge, our work provides the first demonstration that SS4 plays a major role in granule initiation in amyloplasts of cereal grains. The severe defects in granule initiation in the *Tass4* mutant is in contrast to the minor defects in compound starch granule morphology in the rice *Os*SS4b mutant (Toyosawa *et al*., 2016). However, rice has two SS4 paralogs, and the extent to which the other paralog (*Os*SS4a) can compensate for the loss of *Os*SS4b in the endosperm is unknown. Interestingly, *Os*SS4a knockout mutants created by gene-editing were observed to have severe defects in plant growth (Jung *et al*., 2018). Examining endosperm starch in these mutants, as well as in a *osss4a osss4b* double mutant, will be informative of the role of SS4 in a species that already produces compound granules.

### How do *TaSS4* and *Ta*BGC1 control the number of granule initiations?

The increase in initiations per amyloplast that leads to compound granule formation following loss of SS4 in wheat endosperm contrasts the reductions in granule number per chloroplast observed in both Arabidopsis and wheat leaves (Roldán *et al*., 2007)(Fig. 9). Thus, in wheat endosperm, neither *Ta*SS4 nor *Ta*BGC1 is strictly required for the initiation of granules *per se*, but both are required to control the process - such that single A-type granules initiate in amyloplasts during early grain development. It remains to be determined how these proteins exert this control. It is possible that *Ta*SS4 and *Ta*BGC1 together form a single granule initiation per amyloplast, from which the other enzymes of starch biosynthesis can build a single A-type granule (Fig. 11). The formation of this single granule initiation may be enough to suppress the formation of more granules – since the activity of other starch biosynthesis enzymes can be directed towards the growing granule. However, in the absence of this single granule initiation, the other enzymes may start elongating any available substrate, such as soluble maltooligosaccharides, leading to an uncontrolled number of granules being initiated. These enzymes may include starch synthases and starch phosphorylase, which can all elongate maltooligosaccharides in vitro (Hwang *et al*., 2010; Brust *et al*., 2013; Cuesta-Seijo *et al*., 2016). The heterogeneity in granule number among amyloplasts in the endosperm of *Tass4* and *Tabgc1* mutants may reflect stochasticity in the number of initiations per amyloplast that occur in the absence of SS4 or BGC1. It is also possible that some amyloplasts fail to initiate starch granules, but it is very difficult to distinguish empty amyloplasts from other membranous structures in TEM images of the endosperm.

**Fig. 11.**
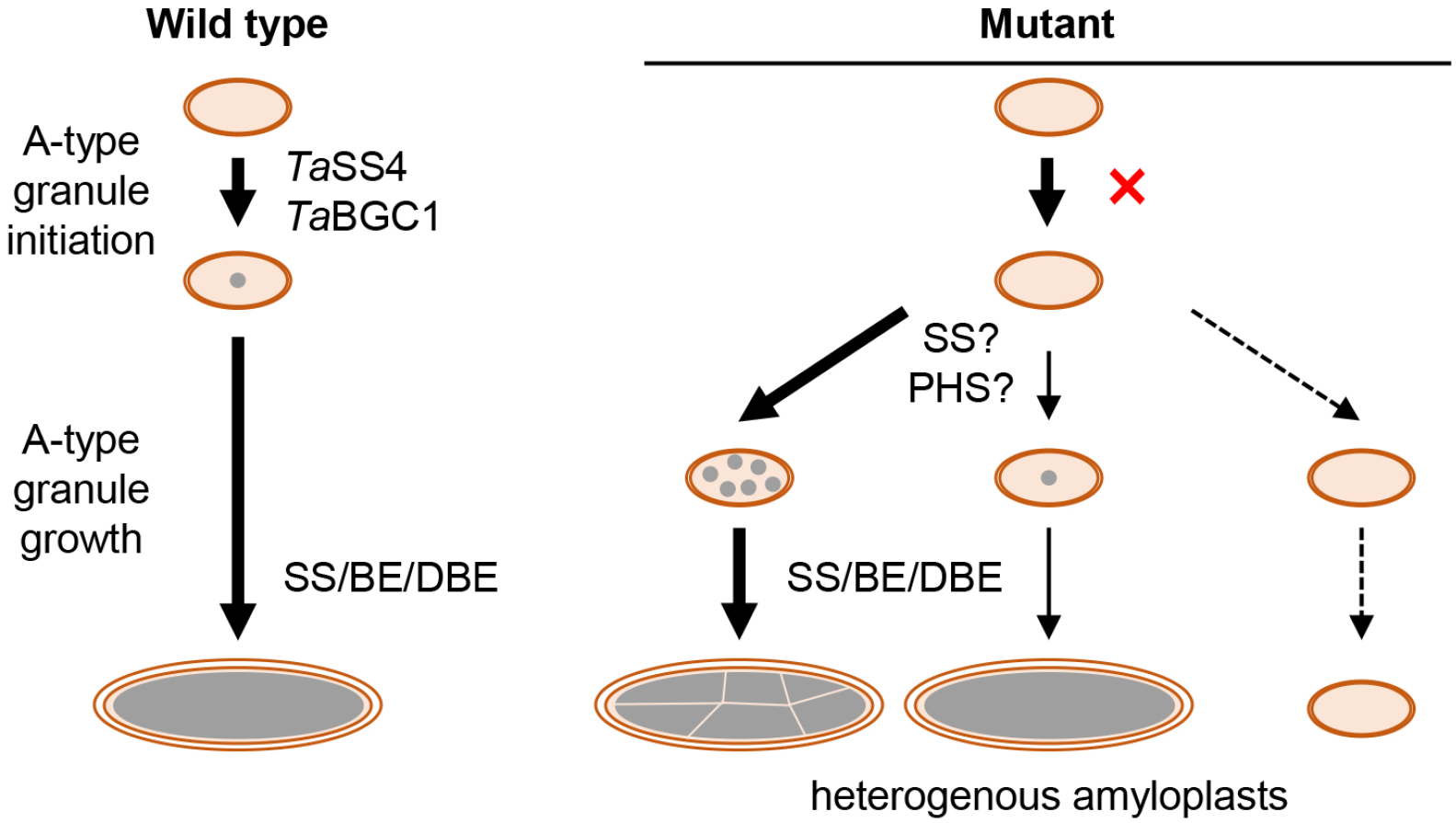
Model of *Ta*SS4 action in endosperm starch initiation. *Ta*SS4 and *Ta*BGC1 are required for the control of normal A-type granule initiation. We propose that they establish a single granule initial that serves as the preferred substrate of other biosynthesis enzymes for building an A-type granule. In the absence of the granule initial, the other biosynthesis enzymes begin to elongate other available substrates such as soluble maltooligosaccharides, which results in the initiation of an undefined number of granules. This leads to heterogeneity among amyloplasts, where most have multiple initiations (leading to a compound granule) and some have normal A-type granules. It is possible that some amyloplasts do not initiate any starch granule, but the prevalence of this is unknown. Abbreviations are SS – starch synthase, PHS – starch phosphorylase, BE – branching enzyme, DBE – debranching enzyme.

It is unknown which features of *Ta*SS4 allow it to initiate a single granule per amyloplast. Notably, distinct patterns of protein localisation have been observed for *At*SS4 in Arabidopsis leaves, and for *Os*SS4b in rice amyloplasts - where it locates to the septum-like structures of compound granules (Toyosawa *et al*., 2016). We are currently exploring the localisation of *Ta*SS4 in amyloplasts of developing grains and whether that could explain a single point of A-type granule initiation. Since granule initiation proteins in Arabidopsis leaves act via protein-protein interactions, searching for interacting proteins may also provide insight on how *Ta*SS4 and *Ta*BGC1 act in wheat endosperm. *At*SS4 is proposed to interact with *At*PTST2 in Arabidopsis leaves (Seung *et al*., 2017). Although we attempted multiple co-immunoprecipitation and pulldown approaches, we failed to find any evidence that *Ta*SS4 and *Ta*BGC1 interact in the endosperm (data not shown). Further work is required to determine if they interact only weakly or transiently. Possible interactions of these proteins with ISOAMYLASE 1 (ISA1) should also be investigated since ISA1 is reported to interact with PTST2 (FLO6) in rice (Peng *et al*., 2014). Notably, *isa1* mutants of barley contain compound granules that resemble those of the *Tass4* mutants (Burton *et al*., 2002), providing a strong indication for ISA1 involvement in granule initiation.

The specific role of *Ta*SS4 in B-type granule initiation must also be further explored. Very few normal round B-type granules were observed in mature grains of the *Tass4* mutant (Fig. 3). Also, at 15 dpa, we observed many compound granules in a linear arrangement in the mutant, raising the possibility that they formed in stromules that normally enclose B-type granules (Fig. 6). Interestingly, Chia et al. (2020) reported that reducing gene dosage of *TaBGC1* in hexaploid wheat can almost eliminate B-type granules while retaining normal A-type granule morphology. By contrast, B-type granule volume was not affected in either of the single homoeolog mutants in *TaSS4*, but it is possible that a further reduction in gene dosage is required to see an effect. However, we noted that while *Ta*SS4 protein levels are highest during early grain development and decrease at the later developmental stages, *Ta*BGC1 transcript and protein levels increase and are highest during the period of B-type granule initiation (Fig. 2; Fig. S3). Thus, it is possible that *Ta*BGC1 has a specific role during B-type granule initiation that is independent of *Ta*SS4.

While other members of the Triticeae (e.g., barley and rye) also have A- and B-type granules, most other grasses produce compound granules in the endosperm (Matsushima *et al*., 2013, 2015). The fact that loss of SS4 or BGC1 gives rise to some compound granules in wheat makes it tempting to speculate that differences in the extent and timing of SS4 and/or BGC1 expression between species could determine whether a given species possesses compound granules. However, the difference between compound and other patterns of granule initiation is unlikely to be so simple. Compound granules of rice have complex structural features, including membranes and septum-like structures that separate each constituent granula (Yun & Kawagoe, 2010; Kawagoe, 2013; Toyosawa *et al*., 2016). Thus, the formation of compound granules in rice is likely to involve multiple genes that control starch synthesis and amyloplast morphogenesis.

### TaSS4 is required for proper granule initiation in leaves and pollen

In leaves of the Arabidopsis *Atss4* mutant, over 75% of chloroplasts had no visible starch granule, and the majority of remaining chloroplasts contained one large granule (Roldán *et al*., 2007; Seung *et al*., 2017). Leaves of the *Tass4* mutant had a percentage of starchless chloroplasts that was comparable to the Arabidopsis mutant, but the remaining chloroplasts mostly contained multiple granules (Fig. 9). The reason for this difference between the Arabidopsis and wheat phenotypes is unknown, but could reflect differences in the compensation mechanism following loss of SS4. The few granules present in the Arabidopsis *Atss4* mutant are likely initiated by SS3, since the *Atss3 Atss4* double mutant is almost starchless (Szydlowski *et al*., 2009; Seung *et al*., 2016). Further work is required to determine whether SS3 initiates the starch granules in leaves of the *Tass4* mutant.

The expression of *Ta*SS4, which shares 56% amino acid sequence identity with *At*SS4 (BLAST pairwise alignment), could largely restore the initiation of multiple granules per chloroplast when expressed in the Arabidopsis *Atss4* mutant. However, the exact role of *Ta*SS4 in granule morphogenesis in leaves remains unclear. Starch granules in the *Tass4* mutant were small and round (Fig. 9), but distinct from the large, rounded granules of the *Atss4* mutant (Fig. 10). *Ta*SS4 expression in the *Atss4* mutant resulted in aberrant granule morphology, which was distinct from both the round granules of *Atss4* and the flattened granules of the wild type. These aberrant granule shapes may result from partial complementation by *Ta*SS4 that achieves an ‘intermediate’ morphology between round and flattened, or abnormal function of *Ta*SS4 in Arabidopsis leaves (e.g., due to missing interaction partners or other regulatory factors).

Despite a reduction in gene dosage to 50% in our single homoeolog wheat mutants, we did not observe an effect on granule number in leaf chloroplasts. On first glance, this is in contrast to a previous report that hexaploid wheat mutants deficient in only *TaSS4-1D* have reduced numbers of granules per chloroplast (Guo *et al*., 2017). However, we showed that some hexaploid cultivars, including the reference cultivar Chinese Spring, have a natural polymorphism that leads to a premature stop codon in *TaSS4-1A* (Fig. 1). It is possible that *TaSS4-1B* is the only functional homoeolog in the *TaSS4-1D* mutants of Guo et al. (2017)(in cultivar Jing411), and thus may have a functional gene dosage of only 33%.

*Ta*SS4 also appears to be required for normal starch synthesis in wheat pollen. Publicly available gene expression data for hexaploid wheat suggests that *TaSS4* is expressed in microspores in addition to leaves, stems, roots and grains (Fig. S3b); and most pollen grains from our *Tass4* mutants were starchless (Fig. 8e, S8). In rice, starch synthesis in pollen appears to be essential for viability, as rice *pgm* mutants lacking pollen starch are sterile (Lee *et al*., 2016). Consistent with this, the pollen from the *Tass4* mutant had significantly reduced fertilisation success in cross-fertilisation experiments (Fig. S8b), and the mutants produced fewer grains per spike (Fig. 8d, S8c). These grains likely result from the small proportion of mutant pollen that contains starch. Further work should examine the effects on granule number and morphology in these starch-containing pollen grains.

## ACKNOWLEDGEMENTS

This work was funded through a John Innes Foundation (JIF) Chris J. Leaver Fellowship (to D.S), a Biotechnology and Biological Sciences Research Council (BBSRC, UK) Future Leader Fellowship BB/P010814/1 (to D.S.), a JIF Rotation Ph.D. studentship (to J.C.) and BBSRC Institute Strategic Programme grants BBS/E/J/000PR9790 and BBS/E/J/000PR9799 (to the John Innes Centre). FJW and JAJ were funded through Quadram BBSRC Institute Strategic Programme grant Food Innovation and Health BB/R012512/1 and its constituent projects BBS/E/F/000PR10343, BBS/E/F/000PR10345 and BBS/E/F/000PR10346. We gratefully acknowledge: Prof. Alison Smith (JIC) for critical reading of this manuscript and her valuable advice throughout the course of this work, Dr. Kay Trafford (NIAB) for critical reading of this manuscript and helpful discussions on granule initiation in wheat, Prof. Cristobal Uauy (JIC) and Dr. Philippa Borrill (University of Birmingham) for their invaluable advice on wheat research. We also thank Dr. George Savva (Quadram Institute) for advice on statistical analysis, and James McLaughlin and Olivia Mohr (JIC) for technical assistance. We thank the JIC Horticultural Services for providing growth facilities and maintenance of plant material, JIC Bioimaging for providing access to microscopes, and the JIC Germplasm Resources Unit for providing the wheat TILLING lines. We also thank three anonymous reviewers for their helpful comments that improved this manuscript.

## AUTHOR CONTRIBUTION

EH and DS conceived and led the study, and designed most of the experiments. EH conducted most of the experiments. JC designed and conducted the Arabidopsis complementation experiments. AWL designed and conducted the transcriptomics analyses. JAJ and FW designed and conducted the HPLC-SEC analyses. JEB designed and conducted TEM experiments and performed sectioning. BF designed and conducted the crosses of the wheat TILLING lines. MH designed the analysis of the granule size distribution data. All authors analysed data. EH and DS wrote the paper with contributions from all authors.

## Notes

### Competing Interest Statement

The authors have declared no competing interest.

## REFERENCES

Abt MR, Pfister B, Sharma M, Eicke S, Bürgy L, Neale I, Seung D, Zeeman SC. 2020. STARCH SYNTHASE5, a Noncanonical Starch Synthase-Like Protein, Promotes Starch Granule Initiation in Arabidopsis. The Plant Cell 32: 2543–2565.

Abt MR, Zeeman SC. 2020. Evolutionary innovations in starch metabolism. Current Opinion in Plant Biology 55: 109–117.

Appels R, Eversole K, Stein N, Feuillet C, Keller B, Rogers J, Pozniak CJ, Choulet F, Distelfeld A, Poland J, et al. 2018. Shifting the limits in wheat research and breeding using a fully annotated reference genome. Science 361: eaar7191.

Bechtel DB, Zayas I, Kaleikau L, Pomeranz Y. 1990. Size-distribution of wheat starch granules during endosperm development. Cereal Chemistry 67: 59–63.

Bolger AM, Lohse M, Usadel B. 2014. Trimmomatic: A flexible trimmer for Illumina sequence data. Bioinformatics 30: 2114–2120.

Borrill P, Ramirez-Gonzalez R, Uauy C. 2016. expVIP: a Customizable RNA-seq Data Analysis and Visualization Platform. Plant Physiology 170: 2172–2186.

Brust H, Orzechowski S, Fettke J, Steup M. 2013. Starch synthesizing reactions and paths: in vitro and in vivo studies. Journal of Applied Glycoscience 60: 3–20.

Burton RA, Jenner H, Carrangis L, Fahy B, Fincher GB, Hylton C, Laurie DA, Parker M, Waite D, Van Wegen S, et al. 2002. Starch granule initiation and growth are altered in barley mutants that lack isoamylase activity. Plant Journal 31: 97–112.

Cave R, Seabrook S, Gidley M, Gilbert R. 2009. Characterization of starch by size-exclusion chromatography: The limitations imposed by shear scission. Biomacromolecules 10: 2245–2253.

Chia T, Chirico M, King R, Ramirez-Gonzalez R, Saccomanno B, Seung D, Simmonds J, Trick M, Uauy C, Verhoeven T, et al. 2020. A carbohydrate-binding protein, B-GRANULE CONTENT 1, influences starch granule size distribution in a dose-dependent manner in polyploid wheat. Journal of Experimental Botany 71: 105–115.

Clavijo BJ, Venturini L, Schudoma C, Accinelli GG, Kaithakottil G, Wright J, Borrill P, Kettleborough G, Heavens D, Chapman H, et al. 2017. An improved assembly and annotation of the allohexaploid wheat genome identifies complete families of agronomic genes and provides genomic evidence for chromosomal translocations. Genome Research 27: 885–896.

Crumpton-Taylor M, Grandison S, Png KMY, Bushby AJ, Smith AM. 2012. Control of starch granule numbers in Arabidopsis chloroplasts. Plant Physiology 158: 905–916.

Crumpton-Taylor M, Pike M, Lu KJ, Hylton CM, Feil R, Eicke S, Lunn JE, Zeeman SC, Smith AM. 2013. Starch synthase 4 is essential for coordination of starch granule formation with chloroplast division during Arabidopsis leaf expansion. New Phytologist 200: 1064–1075.

Cuesta-Seijo JA, Nielsen MM, Ruzanski C, Krucewicz K, Beeren SR, Rydhal MG, Yoshimura Y, Striebeck A, Motawia MS, Willats WGT, et al. 2016. In vitro biochemical characterization of all barley endosperm starch synthases. Frontiers in Plant Science 6: 1–17.

Delatte T, Trevisan M, Parker ML, Zeeman SC. 2005. Arabidopsis mutants Atisa1 and Atisa2 have identical phenotypes and lack the same multimeric isoamylase, which influences the branch point distribution of amylopectin during starch synthesis. Plant Journal 41: 815–830.

Delvallé D, Dumez S, Wattebled F, Roldán I, Planchot V, Berbezy P, Colonna P, Vyas D, Chatterjee M, Ball S, et al. 2005. Soluble starch synthase I: a major determinant for the synthesis of amylopectin in Arabidopsis thaliana leaves. Plant Journal 43: 398–412.

Dumez S, Wattebled F, Dauvillee D, Delvalle D, Planchot V, Ball SG, D’Hulst C. 2006. Mutants of Arabidopsis lacking starch branching enzyme II substitute plastidial starch synthesis by cytoplasmic maltose accumulation. Plant Cell 18: 2694–2709.

Fujita N, Yoshida M, Asakura N, Ohdan T, Miyao A, Hirochika H, Nakamura Y. 2006. Function and characterization of starch synthase I using mutants in rice. Plant Physiology 140: 1070–1084.

Fujita N, Yoshida M, Kondo T, Saito K, Utsumi Y, Tokunaga T, Nishi A, Satoh H, Park JH, Jane JL, et al. 2007. Characterization of SSIIIa-deficient mutants of rice: the function of SSIIIa and pleiotropic effects by SSIIIa deficiency in the rice endosperm. Plant Physiology 144: 2009–2023.

Grefen C, Donald N, Hashimoto K, Kudla J, Schumacher K, Blatt MR. 2010. A ubiquitin-10 promoter-based vector set for fluorescent protein tagging facilitates temporal stability and native protein distribution in transient and stable expression studies. Plant Journal 64: 355–365.

Guo H, Liu Y, Li X, Yan Z, Xie Y, Xiong H, Zhao L, Gu J, Zhao S, Liu L. 2017. Novel mutant alleles of the starch synthesis gene TaSSIVb-D result in the reduction of starch granule number per chloroplast in wheat. BMC Genomics 18: 1–10.

Howard T, Rejab NA, Griffiths S, Leigh F, Leverington-Waite M, Simmonds J, Uauy C, Trafford K. 2011. Identification of a major QTL controlling the content of B-type starch granules in Aegilops. Journal of Experimental Botany 62: 2217–2228.

Hwang S-K, Nishi A, Satoh H, Okita TW. 2010. Rice endosperm-specific plastidial alpha-glucan phosphorylase is important for synthesis of short-chain malto-oligosaccharides. Archives of biochemistry and biophysics 495: 82–92.

Irshad, Guo, Zhang, Gu, Zhao, Xie, Xiong, Zhao, Ding, Ma, et al. 2019. EcoTILLING reveals natural allelic variations in starch synthesis key gene TaSSIV and its haplotypes associated with higher thousand grain weight. Genes 10: 307.

Jung YJ, Nogoy FM, Lee SK, Cho YG, Kang KK. 2018. Application of ZFN for site directed mutagenesis of rice SSIVa gene. Biotechnology and Bioprocess Engineering 23: 108–115.

Karimi M, Inzé D, Depicker A. 2002. GATEWAY vectors for Agrobacterium-mediated plant transformation. Trends in plant science 7: 193–195.

Kawagoe Y. 2013. The characteristic polyhedral, sharp-edged shape of compound-type starch granules in rice endosperm is achieved via the septum-like structure of the amyloplast. Journal of Applied Glycoscience 60: 29–36.

Kersey PJ, Allen JE, Allot A, Barba M, Boddu S, Bolt BJ, Carvalho-Silva D, Christensen M, Davis P, Grabmueller C, et al. 2018. Ensembl Genomes 2018: An integrated omics infrastructure for non-vertebrate species. Nucleic Acids Research 46: D802–D808.

Krasileva K V., Vasquez-Gross HA, Howell T, Bailey P, Paraiso F, Clissold L, Simmonds J, Ramirez-Gonzalez RH, Wang X, Borrill P, et al. 2017. Uncovering hidden variation in polyploid wheat. Proceedings of the National Academy of Sciences 114: E913–E921.

Langeveld SMJ, Van wijk R, Stuurman N, Kijne JW, de Pater S. 2000. B-type granule containing protrusions and interconnections between amyloplasts in developing wheat endosperm revealed by transmission electron microscopy and GFP expression. Journal of Experimental Botany 51: 1357–1361.

Lee SK, Eom JS, Hwang SK, Shin D, An G, Okita TW, Jeon JS. 2016. Plastidic phosphoglucomutase and ADP-glucose pyrophosphorylase mutants impair starch synthesis in rice pollen grains and cause male sterility. Journal of Experimental Botany 67: 5557–5569.

Lu K-J, Pfister B, Jenny C, Eicke S, Zeeman SC. 2018. Distinct functions of STARCH SYNTHASE 4 domains in starch granule formation. Plant Physiology 176: 566–581.

Maccaferri M, Harris NS, Twardziok SO, Pasam RK, Gundlach H, Spannagl M, Ormanbekova D, Lux T, Prade VM, Milner SG, et al. 2019. Durum wheat genome highlights past domestication signatures and future improvement targets. Nature Genetics 51: 885–895.

Matsushima R, Maekawa M, Sakamoto W. 2015. Geometrical formation of compound starch grains in rice implements Voronoi diagram. Plant and Cell Physiology 56: 2150–2157.

Matsushima R, Yamashita J, Kariyama S, Enomoto T, Sakamoto W. 2013. A phylogenetic re-evaluation of morphological variations of starch grains among Poaceae species. Journal of Applied Glycoscience 60: 131–135.

Morell MK, Kosar-Hashemi B, Cmiel M, Samuel MS, Chandler P, Rahman S, Buleon A, Batey IL, Li Z. 2003. Barley sex6 mutants lack starch synthase lla activity and contain a starch with novel properties. Plant Journal 34: 173–185.

Parker ML. 1985. The relationship between A-type and B-type starch granules in the developing endosperm of wheat. Journal of Cereal Science 3: 271–278.

Patro R, Duggal G, Love MI, Irizarry RA, Kingsford C. 2017. Salmon provides fast and bias-aware quantification of transcript expression. Nature Methods 14: 417–419.

Peng C, Wang Y, Liu F, Ren Y, Zhou K, Lv J, Zheng M, Zhao S, Zhang L, Wang C, et al. 2014. FLOURY ENDOSPERM6 encodes a CBM48 domain-containing protein involved in compound granule formation and starch synthesis in rice endosperm. Plant Journal 77: 917–930.

Ragel P, Streb S, Feil R, Sahrawy M, Annunziata MG, Lunn JE, Zeeman S, Mérida Á. 2013. Loss of starch granule initiation has a deleterious effect on the growth of Arabidopsis plants due to an accumulation of ADP-glucose. Plant Physiology 163: 75–85.

Roldán I, Wattebled F, Mercedes Lucas M, Delvallé D, Planchot V, Jiménez S, Pérez R, Ball S, D’Hulst C, Mérida A. 2007. The phenotype of soluble starch synthase IV defective mutants of Arabidopsis thaliana suggests a novel function of elongation enzymes in the control of starch granule formation. Plant Journal 49: 492–504.

Saito M, Tanaka T, Sato K, Vrinten P, Nakamura T. 2017. A single nucleotide polymorphism in the “Fra” gene results in fractured starch granules in barley. Theoretical and Applied Genetics 131: 353–364.

Seung D. 2020. Amylose in starch: towards an understanding of biosynthesis, structure and function. New Phytologist In Press: nph.16858.

Seung D, Boudet J, Monroe JD, Schreier TB, David LC, Abt M, Lu K-J, Zanella M, Zeeman SC. 2017. Homologs of PROTEIN TARGETING TO STARCH control starch granule initiation in Arabidopsis leaves. Plant Cell 29: 1657–1677.

Seung D, Lu K-J, Stettler M, Streb S, Zeeman SC. 2016. Degradation of glucan primers in the absence of Starch Synthase 4 disrupts starch granule initiation in Arabidopsis. Journal of Biological Chemistry 291: 20718–20728.

Seung D, Schreier TB, Bürgy L, Eicke S, Zeeman SC. 2018. Two plastidial coiled-coil proteins are essential for normal starch granule initiation in Arabidopsis. Plant Cell 30: 1523–1542.

Seung D, Smith A. 2019. Starch granule initiation and morphogenesis–progress in Arabidopsis and cereals. Journal of Experimental Botany 70: 771–784.

Seung D, Soyk S, Coiro M, Maier BA, Eicke S, Zeeman SC. 2015. PROTEIN TARGETING TO STARCH is required for localising GRANULE-BOUND STARCH SYNTHASE to starch granules and for normal amylose synthesis in Arabidopsis. PLOS Biology 13: e1002080.

Smith AM, Zeeman SC. 2020. Starch: A flexible, adaptable carbon store coupled to plant growth. Annual Review of Plant Biology 71: 217–245.

Suh DS, Verhoeven T, Denyer K, Jane JL. 2004. Characterization of Nubet and Franubet barley starches. Carbohydrate Polymers 56: 85–93.

Sundberg M, Pfister B, Fulton D, Bischof S, Delatte T, Eicke S, Stettler M, Smith SM, Streb S, Zeeman SC. 2013. The heteromultimeric debranching enzyme involved in starch synthesis in Arabidopsis requires both Isoamylase1 and Isoamylase2 subunits for complex stability and activity. PLOS One 8: e75223.

Szydlowski N, Ragel P, Hennen-Bierwagen T a, Planchot V, Myers AM, Mérida A, D’Hulst C, Wattebled F. 2011. Integrated functions among multiple starch synthases determine both amylopectin chain length and branch linkage location in Arabidopsis leaf starch. Journal of Experimental Botany: 1–13.

Szydlowski N, Ragel P, Raynaud S, Lucas MM, Roldán I, Montero M, Muñoz FJ, Ovecka M, Bahaji A, Planchot V, et al. 2009. Starch granule initiation in Arabidopsis requires the presence of either class IV or class III starch synthases. Plant Cell 21: 2443–2457.

Tanaka E, Ral JPF, Li S, Gaire R, Cavanagh CR, Cullis BR, Whan A. 2017. Increased accuracy of starch granule type quantification using mixture distributions. Plant Methods: 1–7.

Toyosawa Y, Kawagoe Y, Matsushima R, Crofts N, Ogawa M, Fukuda M, Kumamaru T, Okazaki Y, Kusano M, Saito K, et al. 2016. Deficiency of starch synthase IIIa and IVb alters starch granule morphology from polyhedral to spherical in rice endosperm. Plant Physiology 170: 1255–1270.

Tuncel A, Corbin KR, Ahn-Jarvis J, Harris S, Hawkins E, Smedley MA, Harwood W, Warren FJ, Patron NJ, Smith AM. 2019. Cas9-mediated mutagenesis of potato starch-branching enzymes generates a range of tuber starch phenotypes. Plant Biotechnology Journal 17: 2259–2271.

Vandromme C, Spriet C, Dauvillée D, Courseaux A, Putaux J, Wychowski A, Krzewinski F, Facon M, D’Hulst C, Wattebled F. 2019. PII1: a protein involved in starch initiation that determines granule number and size in Arabidopsis chloroplast. New Phytologist 221: 356–370.

Vilaplana F, Hasjim J, Gilbert RG. 2012. Amylose content in starches: Toward optimal definition and validating experimental methods. Carbohydrate Polymers 88: 103–111.

Wang YJ, White P, Pollak L, Jane J. 1993. Characterization of starch structures of 17 maize endosperm mutant genotypes with Oh43 inbred line background. Cereal Chemistry 70: 171–179.

Wu AC, Li E, Gilbert RG. 2014. Exploring extraction/dissolution procedures for analysis of starch chain-length distributions. Carbohydrate Polymers 114: 36–42.

Xiang D, Quilichini TD, Liu Z, Gao P, Pan Y, Li Q, Nilsen KT, Venglat P, Esteban E, Pasha A, et al. 2019. The transcriptional landscape of polyploid wheats and their diploid ancestors during embryogenesis and grain development[OPEN]. Plant Cell 31: 2888–2911.

Yun MS, Kawagoe Y. 2010. Septum formation in amyloplasts produces compound granules in the rice endosperm and is regulated by plastid division proteins. Plant and Cell Physiology 51: 1469–1479.

Zhang X, Henriques R, Lin S-S, Niu Q-W, Chua N-H. 2006. Agrobacterium-mediated transformation of Arabidopsis thaliana using the floral dip method. Nature protocols 1: 641–6.

Zhang X, Myers A, James M. 2005. Mutations affecting starch synthase III in Arabidopsis alter leaf starch structure and increase the rate of starch synthesis. Plant Physiology 138: 663–674.

Zhang X, Szydlowski N, Delvallé D, D’Hulst C, James MG, Myers AM. 2008. Overlapping functions of the starch synthases SSII and SSIII in amylopectin biosynthesis in Arabidopsis. BMC Plant Biology 8: 96.

