## Supplemental Figures and Tables for "STARCH SYNTHASE 4 is required for normal starch granule initiation in amyloplasts of wheat endosperm"

**Fig. S1** Amino acid sequence alignment of the wheat *Ta*SS4 homoeologs. **(a)** The three homoeologs (1A, 1B and 1D) in the hexaploid cultivar Chinese Spring (CS), and the two homoeologs (1A and 1B) from the tetraploid cultivar Kronos, were aligned using the ClustalO program. Residues are coloured according to side-chain properties: acidic (blue), basic (magenta), hydrophobic (red), or other (green). Asterisks (*) under the alignment indicate identical residues, while colon (:) and period (.) indicate strongly or weakly similar side-chain properties, respectively. The premature stop codon in the *TaSS4-1A* homoeolog of Chinese Spring is highlighted in red. **(b)** Percentage identity values between sequences in **(a)**.

## (a)

```
CS_TraesCS1A02G353300.2         MACSAAAGVEATALLSPRCPAPSPPDGRSRRRLALASGTRHRSLRAAAQRPHKSATGADP   60
Kronos_TraesCS1A02G353300.2     MACSAAAGVEATALLSPRCPAPSPPDGRSRRRLALASGTRHRSLRAAAQRPHKSATGADP   60
CS_TraesCS1B02G368500.1         MACSAAAGVEATALLSPRCPAPSPPDGRSRRRLALASRTRHRSLRAAAQRPHKSTTGADP   60
Kronos_TraesCS1B02G368500.1     MACSAAAGVEATALLSPRCPAPSPPDGRSRRRLALASRTRHRSLRAAAQRPHKSTTGADP   60
CS_TraesCS1D02G356900.1         MACSAAAGVEATALLSPRCPAPSPPDGRSRRRLALASRTRHRSLRAAAQRPHKSTTGADP   60
                                ************************************* ****************:*****

CS_TraesCS1A02G353300.2         LYNNRANVRSDEASVSAEKERQRKYNDGDGISNLKLEDLVGMIQNTEKNILLLNQARLQA   120
Kronos_TraesCS1A02G353300.2     LYNNRANVRSDEASVSAEKERQRNYNDGDGISNLKLEDLVGMIQNTEKNILLLNQARLQA   120
CS_TraesCS1B02G368500.1         L-NNRANVRSDEAAVSAEKERQRKYNDGDGISNLQLEDLVGMIQNTEKNILLLNQARLQA   119
Kronos_TraesCS1B02G368500.1     L-NNRANVRSDEAAVSAEKERQRKYNDGDGISNLQLEDLVGMIQNTEKNILLLNQARLQA   119
CS_TraesCS1D02G356900.1         L-NNRANPRSDEAAVSAEKERQRKYNDGDGISNLQLEDLVGLIQNTEKNILLLNQARLQA   119
                                * ***** *****:*********:**********:******:******************

CS_TraesCS1A02G353300.2         MEHADKVLKEKEALQRKINILETRLSETDEQHKLSSEGNFSDSPLALELGILKEENILLK   180
Kronos_TraesCS1A02G353300.2     MEHADKVLKEKEALQRKINILETRLSETDEQHKLSSEGNFSDSPLALELGILKEENILLK   180
CS_TraesCS1B02G368500.1         MEHADKILKEKEALQRKINILETRLSEIDSQHKLSSEGNFSDSPLALEFDVLKEENIVLK   179
Kronos_TraesCS1B02G368500.1     MEHADKILKEKEALQRKINILETRLSEIDSQHKLSSEGNFSDSPLALEFDVLKEENIVLK   179
CS_TraesCS1D02G356900.1         MEHADKILKEKEALQRKINILETRLSETNAQHKLSSEGNVSDSPLLLEFDVLKEDNILLK   179
                                ******:******************** : *********.***** **:.:***:**:**

CS_TraesCS1A02G353300.2         EDI*FFKTKLIEVAEIEEGIFKLEKERALLDASLRELESRFIAAQADTMKLGPRDAWWEK   239
Kronos_TraesCS1A02G353300.2     EDIEFFKTKLIEVAEIEEGIFKLEKERALLDASLRELESRFIAAQADTMKLGPRDAWWEK   240
CS_TraesCS1B02G368500.1         EDIEFFKTKLIEVAETEEGIFKLEKERALLDASLRELESRFIAAQANMMKLGPRDAWWEK   239
Kronos_TraesCS1B02G368500.1     EDIEFFKTKLIEVAETEEGIFKLEKERALLDASLRELESRFIAAQANMMKLGPRDAWWEK   239
CS_TraesCS1D02G356900.1         EDIEFFKTKLIEIAETEEGIFKLEKERALLDASLRELESRFIAAQANMMKLGPRDAWWEK   239
                                *** ********:** ******************************: ************

CS_TraesCS1A02G353300.2         VEKLEDLLETTANQVEHAAVILDHNHDLQDRLDKLEASLQAANISKFSCSLVDILQQKVK   299
Kronos_TraesCS1A02G353300.2     VEKLEDLLETTANQVEHAAVILDHNHDLQDRLDKLEASLQAANISKFSCSLVDILQQKVK   300
CS_TraesCS1B02G368500.1         VEKLEDLLETTANQVEHAAVILDRNHDLQDRLEKLEASLQAANISKFSCSLVDLLQQKVK   299
Kronos_TraesCS1B02G368500.1     VEKLEDLLETTANQVEHAAVILDRNHDLQDRLEKLEASLQAANISKFSCSLVDLLQQKVK   299
CS_TraesCS1D02G356900.1         VEKLEDLLETTANQVEHAAVILDHNHDLQDRLDNLEASLQAANISKFSCSLVDLLQQKVK   299
                                ***********************:********::********************:*****

CS_TraesCS1A02G353300.2         LVEERFQACNCEMHSQIELYEHSIVEFHDTLSKLIEESEKRSLENFTGNMPSELWSKISL   359
Kronos_TraesCS1A02G353300.2     LVEECFQACNCEMHSQIELYEHSIVEFHDTLSKLIEESEKRSLENFTGNMPSELWSKISL   360
CS_TraesCS1B02G368500.1         LVEERFQACNREMHSQIELYEHSIVEFHDTLSKLIEESEKRSLENFTGNMPSELWSKISL   359
Kronos_TraesCS1B02G368500.1     LVEERFQACNREMHSQIELYEHSIVEFHDTLSKLIEESEKRSLENFTGNMPSELWSKISL   359
CS_TraesCS1D02G356900.1         LVEDRFQACNSEMHSQIELYEHSIVEFHDTLSKLIEESEKRSLENFTGNMPSELWSKISL   359
                                ***: ***** *************************************************

CS_TraesCS1A02G353300.2         LTDGWLLEKKISYNDASMLREMVHKRDSRLREAYLSYRGTENREVMDNLLKMALPGTSSG   419
Kronos_TraesCS1A02G353300.2     LTDGWLLEKKISYNDASMLREMVHKRDSRLREAYLSYRGTENREVMDNLLKMALPGTSSG   420
CS_TraesCS1B02G368500.1         LTDGWLLEKKISYSDASMLREMVQKRDNRLREAYLSYRGTENREVMDNLLKMALPGTSSG   419
Kronos_TraesCS1B02G368500.1     LTDGWLLEKKISYSDASMLREMVQKRDNRLREAYLSYRGTENREVMDNLLKMALPGTSSG   419
CS_TraesCS1D02G356900.1         LIDGWLLEKKIAYNDASMLREMVRKRDSRLREAYLSYRGTENRDVMDSFLKMALPGTSSG   419
                                * *********:*.*********:***.***************:***.:***********
```

```
CS_TraesCS1A02G353300.2         LHIAHIAAEMAPVAKVGGLADVISGLGKALQKKGHLVEIILPKYDCMQVDQVSNLRVLDV   479
Kronos_TraesCS1A02G353300.2     LHIAHIAAEMAPVAKVGGLADVISGLGKALQKKGHLVEIILPKYDCMQVDQVSNLRVLDV   480
CS_TraesCS1B02G368500.1         LHIAHIAAEMAPVAKVGGLADVISGLGKALQKKGHLVEIILPKYDCMQVDQVSNLRVLDV   479
Kronos_TraesCS1B02G368500.1     LHIAHIAAEMAPVAKVGGLADVISGLGKALQKKGHLVEIILPKYDCMQVDQVSNLRVLDV   479
CS_TraesCS1D02G356900.1         LHIAHIAAEMAPVAKVGGLADVISGLGKALQKKGHLVEIILPKYDCMQVDQVSNLKVLDV   479
                                *******************************************************:****

CS_TraesCS1A02G353300.2         LVQSYFEGNMFNNKIWTGTVEGLPVYFIEPQHPAMFFSRAQYYGEHDDFKRFSYFSRAAL   539
Kronos_TraesCS1A02G353300.2     LVQSYFEGNMFNNKIWTGTVEGLPVYFIEPQHPAMFFSRAQYYGEHDDFKRFSYFSRAAL   540
CS_TraesCS1B02G368500.1         LVQSYFEGNMFNNKIWTGTVEGLPVYFIEPQHPAMFFSRAHYYGEHDDFKRFSYFSRAAL   539
Kronos_TraesCS1B02G368500.1     LVQSYFEGNMFNNKIWTGTVEGLPVYFIEPQHPAMFFSRAHYYGEHDDFKRFSYFSRAAL   539
CS_TraesCS1D02G356900.1         LVQSYFEGNMFNNKIWTGTVEGLPVYFIEPQHPAMFFSRAQYYGEHDDFKRFTYFSRAAL   539
                                ****************************************:***********:*******

CS_TraesCS1A02G353300.2         ELLYQSGKKVDIIHCHDWQTAFVAPLYWDVYANLGFNSARICFTCHNFEYQGTAPARDLA   599
Kronos_TraesCS1A02G353300.2     ELLYQSGKKVDIIHCHDWQTAFVAPLYWDVYANLGFNSARICFTCHNFEYQGTAPARDLA   600
CS_TraesCS1B02G368500.1         ELLYQSGKKVDIIHCHDWQTAFVAPLYWDVYANLGFNSARICFTCHNFEYQGTAPARDLA   599
Kronos_TraesCS1B02G368500.1     ELLYQSGKKVDIIHCHDWQTAFVAPLYWDVYANLGFNSARICFTCHNFEYQGTAPARDLA   599
CS_TraesCS1D02G356900.1         ELLYQSGKKVDIIHCHDWQTAFVAPLYWDVYANLGFNSARICFTCHNFEYQGTAPARDLA   599
                                ************************************************************

CS_TraesCS1A02G353300.2         WCGLDVEHLDRPDRMRDNSHGRINAVKGAVVYSNIVTTVSPTYALEVRSEGGRGLQDTLK   659
Kronos_TraesCS1A02G353300.2     WCGLDVEHLDRPDRMRDNSHGRINAVKGAVVYSNIVTTVSPTYALEVRSEGGRGLQDTLK   660
CS_TraesCS1B02G368500.1         WCGLDVEHLDRPDRMRDNSHGRINAVKGAVVYSNIVTTVSPTYALEVRSEGGRGLQDTLK   659
Kronos_TraesCS1B02G368500.1     WCGLDVEHLDRPDRMRDNSHGRINAVKGAVVYSNIVTTVSPTYALEVRSEGGRGLQDTLK   659
CS_TraesCS1D02G356900.1         WCGLDVEHLDRPDRMRDNSHGRINAVKGAIVYSNIVTTVSPTYALEVRSEGGRGLQDTLK   659
                                *****************************:******************************

CS_TraesCS1A02G353300.2         VHSRKFLGILNGIDTDTWNPSTDRYLKVQYNAKDLQGKAANKAALREQLNLASAYPSQPL   719
Kronos_TraesCS1A02G353300.2     VHSRKFLGILNGIDTDTWNPSTDRYLKVQYNAKDLQGKAANKAALREQLNLASAYPSQPL   720
CS_TraesCS1B02G368500.1         VHSRKFLGILNGIDTDTWNPCTDRYLKVQYNAKDLQGKAANKAALREQLNLASAYPSQPL   719
Kronos_TraesCS1B02G368500.1     VHSRKFLGILNGIDTDTWNPCTDRYLKVQYNAKDLQGKAANKAALREQLNLASAYPSQPL   719
CS_TraesCS1D02G356900.1         VHSRKFLGILNGIDTDTWNPSTDRYLKVQYNAKDLQGKAANKAALREQLNLASAYPSQPL   719
                                ********************.***************************************

CS_TraesCS1A02G353300.2         VGCITRLVAQKGVHLIRHAIYKTAELGGQFVLLGSSPVPEIQREFEGIADHFQNNNNIRL   779
Kronos_TraesCS1A02G353300.2     VGCITRLVAQKGVHLIRHAIYKTAELGGQFVLLGSSPVPEIQREFEGIADHFQNNNNIRL   780
CS_TraesCS1B02G368500.1         VGCITRLVAQKGVHLIRHAIYKTAELGGQFVLLGSSPVPEIQREFEGIADHFQNNNNIRL   779
Kronos_TraesCS1B02G368500.1     VGCITRLVAQKGVHLIRHAIYKTAELGGQFVLLGSSPVPEIQREFEGIADHFQNNNNIRL   779
CS_TraesCS1D02G356900.1         VGCITRLVAQKGVHLIRHAIYKTAELGGQFVLLGSSPVPEIQREFEGIADHFQNNNNIRL   779
                                ************************************************************

CS_TraesCS1A02G353300.2         ILKYDDALSHCIYAASDMFIVPSIFEPCGLTQMIAMRYGSVPIVRKTGGLNDSVFDFDDE   839
Kronos_TraesCS1A02G353300.2     ILKYDDALSHCIYAASDMFIVPSIFEPCGLTQMIAMRYGSVPIVRKTGGLNDSVFDFDDE   840
CS_TraesCS1B02G368500.1         ILKYDDALSHCIYAASDMFIVPSIFEPCGLTQMIAMRYGSVPIVRKTGGLNDSVFDFDDE   839
Kronos_TraesCS1B02G368500.1     ILKYDDALSHCIYAASDMFIVPSIFEPCGLTQMIAMRYGSVPIVRKTGGLNDSVFDFDDE   839
CS_TraesCS1D02G356900.1         ILKYDDALSHCIYAASDMFIVPSIFEPCGLTQMIAMRYGSVPIVRKTGGLNDSVFDFDDE   839
                                ************************************************************

CS_TraesCS1A02G353300.2         TIPMEVRNGFTFVKADEQGLSSAMERAFNCYTRKPEVWKQLVQKDMTIDFSWDTSASQYE   899
Kronos_TraesCS1A02G353300.2     TIPMEVRNGFTFVKADEQGLSSAMERAFNCYTRKPEVWKQLVQKDMTIDFSWDTSASQYE   900
CS_TraesCS1B02G368500.1         TIPMEVRNGFTFVKADEQGLSSAMERAFNCYTRKPEVWKQLVQKDMTIDFSWDTSASQYE   899
Kronos_TraesCS1B02G368500.1     TIPMEVRNGFTFVKADEQGLSSAMERAFNCYTRKPEVWKQLVQKDMTIDFSWDTSASQYE   899
CS_TraesCS1D02G356900.1         TIPMEVRNGFTFVKADEQGLSSAMERAFNCYTRKPEVWKQLVQKDMTIDFSWDTSASQYE   899
                                ************************************************************

CS_TraesCS1A02G353300.2         DIYQKAVARARAVA 913
Kronos_TraesCS1A02G353300.2     DIYQKAVARARAVA 914
CS_TraesCS1B02G368500.1         DIYQKAVARARAVA 913
Kronos_TraesCS1B02G368500.1     DIYQKAVARARAVA 913
CS_TraesCS1D02G356900.1         DIYQKAVARARAVA 913
                                **************
```

**(b)**

| | CS_TraesCS**1A**02G353300.2 | Kronos_TraesCS**1A**02G353300.2 | CS_TraesCS**1B**02G368500.1 | Kronos_TraesCS**1B**02G368500.1 | CS_TraesCS**1D**02G356900.1 |
|---|---|---|---|---|---|
| CS_TraesCS**1A**02G353300.2 | | | | | |
| Kronos_TraesCS**1A**02G353300.2 | 99 | | | | |
| CS_TraesCS**1B**02G368500.1 | 97 | 97 | | | |
| Kronos_TraesCS**1B**02G368500.1 | 97 | 97 | 100 | | |
| CS_TraesCS**1D**02G356900.1 | 96 | 96 | 97 | 97 | |

**Fig. S2** *Ta*SS4 and *Ta*BGC1 antibodies can detect both A- and B-genome homoeologs. **(a)** *Ta*SS4-1A and *Ta*SS4-1B proteins were transiently expressed in *Nicotiana benthamiana* leaves under the CaMV 35S promoter and with a C-terminal HA-tag. Three replicate protein extracts were prepared from different leaves (numbered 1-3), and were immunoblotted with the anti-*Ta*SS4 (upper panel) or anti-HA (lower panel) antibodies. Lanes were loaded on an equal fresh weight basis. The migration of molecular weight markers is indicated in kilodaltons (kDa) to the left of each panel. **(b)** As for **(a)**, but with *Ta*BGC1-4A and *Ta*BGC1-4B, immunoblotted with the anti-*Ta*BGC1 (upper panel) or anti-HA (lower panel) antibodies.

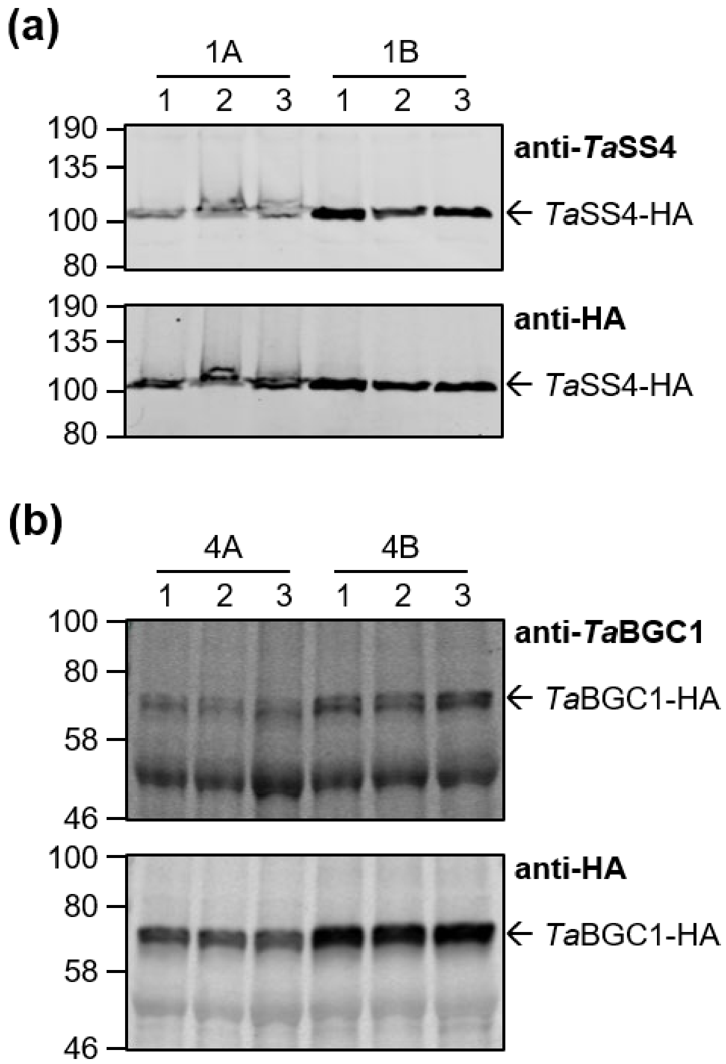

**Fig. S3** Expression analyses of *TaSS4* and *TaBGC1*. **(a)** Transcript levels of *TaSS4* and *TaBGC1* during grain development in tetraploid wheat. Whole developing caryopses at 11, 16 and 21 dpa and endosperm samples at 8 and 22 dpa were determined in transcripts per million (TPM). Transcript levels for both homoeologs were combined to give a single TPM value for each gene. Fold change was calculated between 11 and 16 dpa (caryopsis), 11 and 21 dpa (caryopsis), and 8 and 22 dpa (endosperm). Raw data for caryopsis samples (*n*=3 for each stage) were extracted from Maccaferri *et al.* (2019) and Xiang *et al.* (2019) for endosperm samples (*n*=2 for each stage). **(b)** Expression levels of *TaSS4* homoeologs in different organs of hexaploid wheat. Data were retrieved from the wheat expression browser (http://www.wheat-expression.com; Borrill et al. 2016) using the CS_grain, CS_development and microspores datasets.

**(a)**

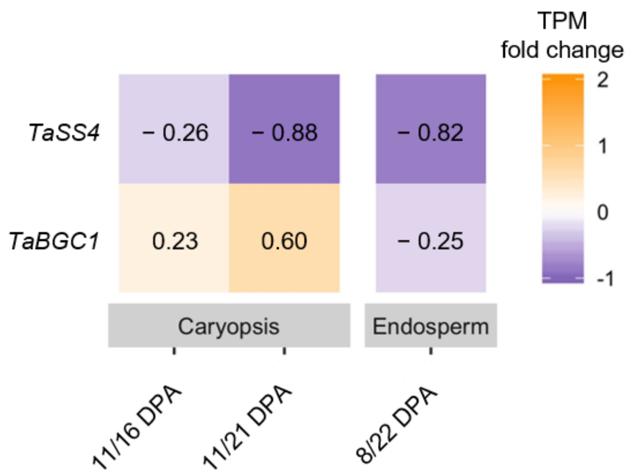


**(b)**

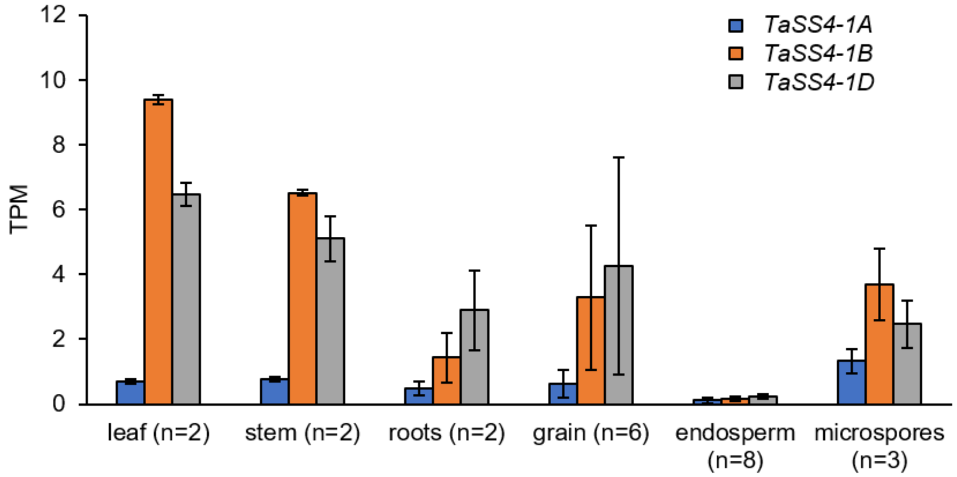

**Fig. S4** Granule morphology of *Tass4-1* after backcrossing and *Tass4-2*. **(a)** Scanning Electron Microscopy (SEM) images of endosperm starch from mature grains of backcrossed *Tass4-1* lines. The negative segregant control (AA BB) and double mutant (*aa bb*) were re-isolated in the F2 generation after backcrossing twice to the wild type ($BC_2F_2$). Bars = 20 µm. **(b)** Size distribution of endosperm starch granules from backcrossed and non-backcrossed *Tass4-1* mutants quantified using a Coulter counter. Values represent mean (solid line) ± SEM (shading) of *n*=4-7 replicate starch extractions, each from grains harvested from a separate plant. **(c)** As for (a), but with *Tass4-2* lines - including the negative segregant control (AA BB), single *TaSS4-1B* mutant (AA *bb*) and double mutant (*aa bb*).

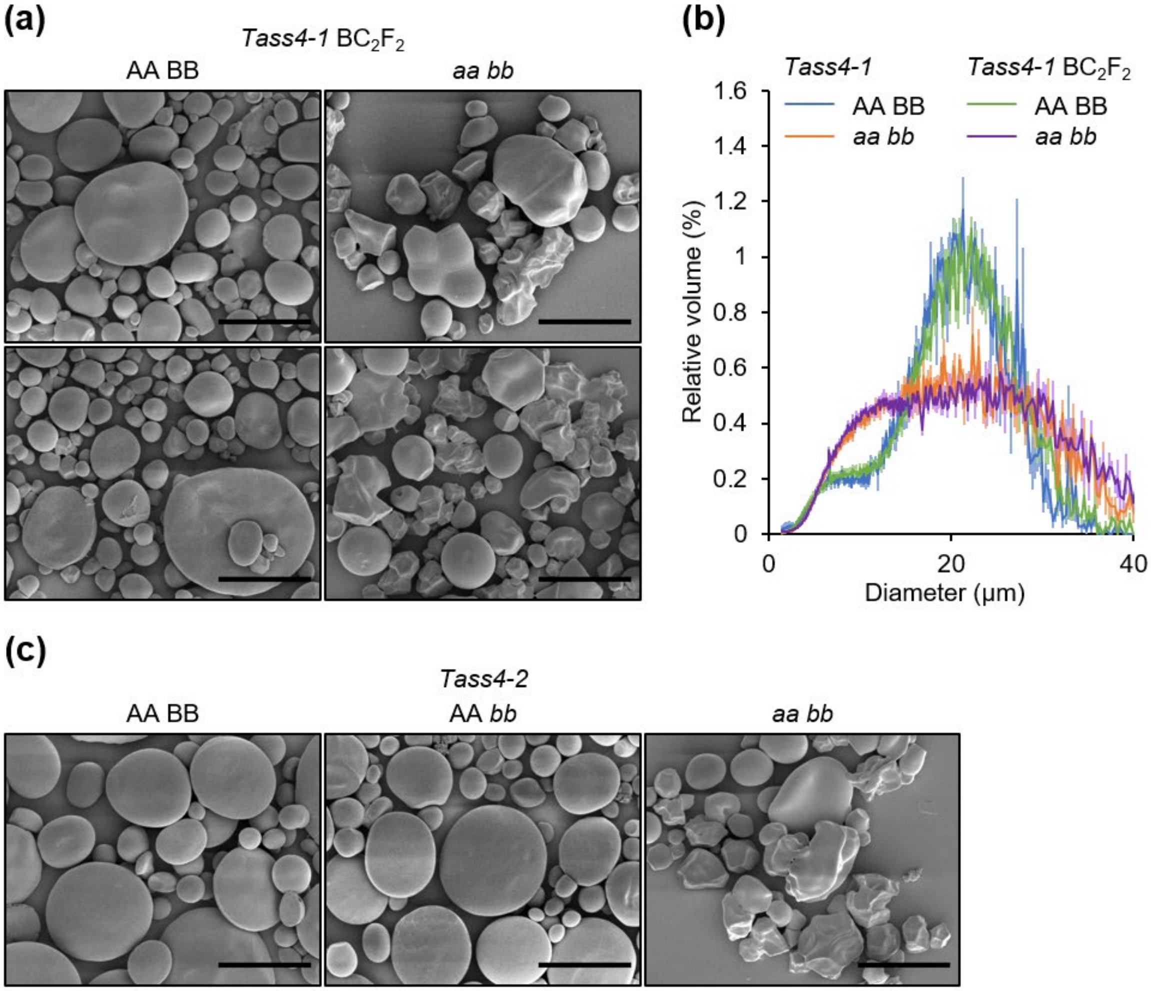

**Fig. S5** Average chain length distributions of debranched *Tass4-1* starch. Starch was purified from whole grains and debranched with isoamylase prior to analysis using SEC-HPLC. The y-axis, w(Log$V_h$) is expressed as the weight distribution based on the relationship between elution volume and hydrodynamic radius (Log$V_h$) for pullulan standards. Chains with DP>100 represent amylose chains and were used to calculate amylose content. Each line represents the average of three different plants.

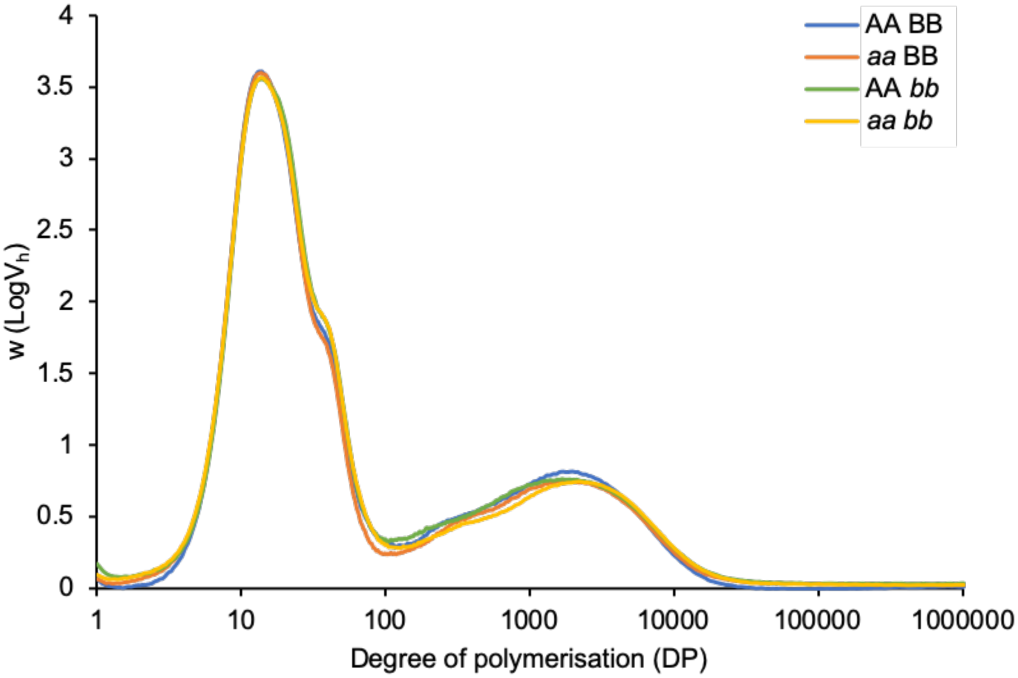

**Fig. S6** Starch from developing grains of *Tabgc1*. Endosperm starch from developing grains at 8, 15 and 20 dpa were observed using scanning electron microscopy (SEM). Bars = 20 µm.

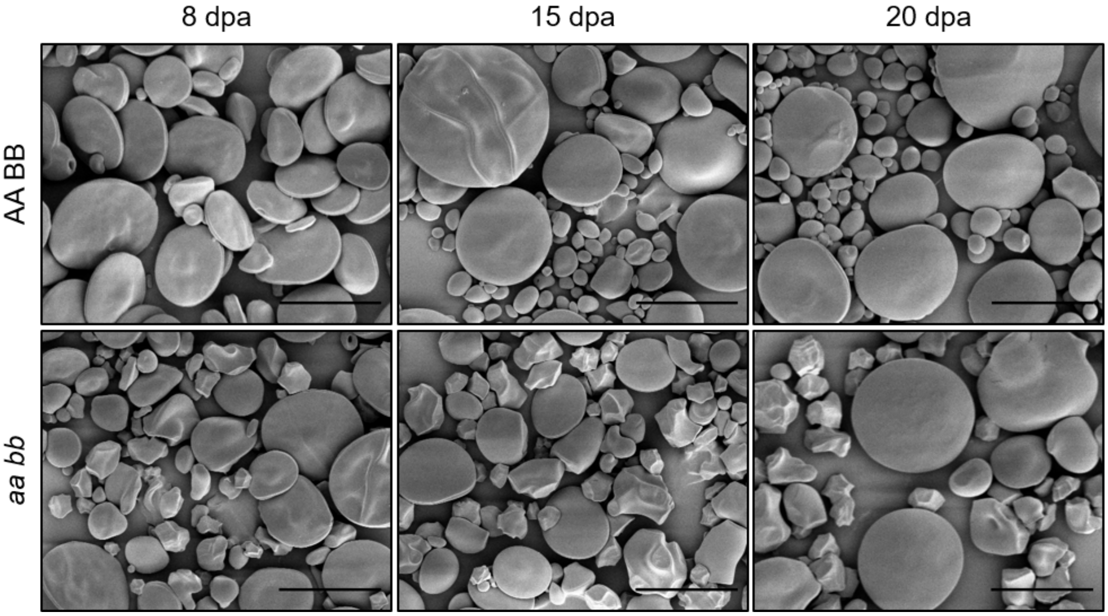

**Fig. S7** Average grain weight of *Tass4* plants. The average weight of individual grains at maturity was calculated for *Tass4-1* [backcrossed ($BC_2F_2$) and non-backcrossed] and *Tass4-2* plants. Each black data point represents the value for an individual plant, while the red data point represents the mean (*n*=2-7 plants per genotype). One-way ANOVA and Tukey's posthoc test revealed no significant differences between the genotypes.

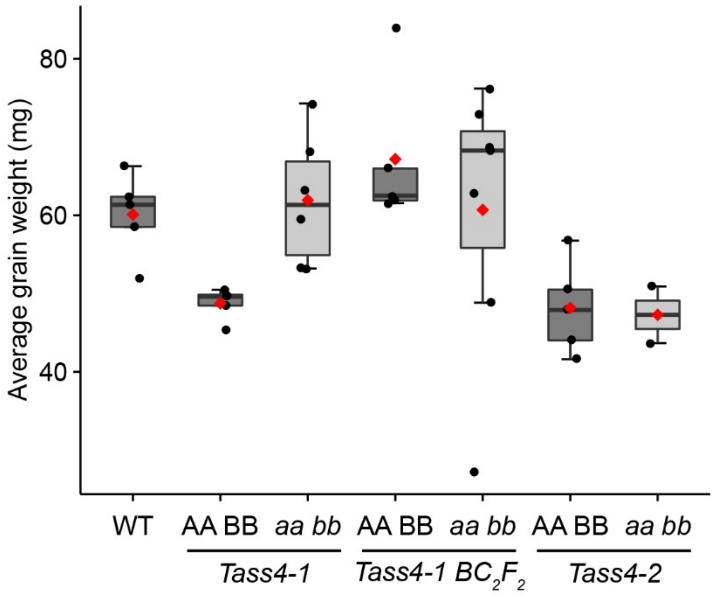

**Fig. S8** Starch in pollen grains of *Tass4* mutants. **(a)** Scoring of starch-containing and starchless pollen in *Tass4-1* [backcrossed ($BC_2F_2$) and non-backcrossed] and *Tass4-2*. Mature pollen grains were harvested from whole anthers and stained with iodine solution for observation with light microscopy. Pollen was scored as starch-containing or starchless (*n*=99-134 pollen grains per plant, *n*=2-7 plants per genotype). Different letters indicate significant differences at $p<0.05$ under a one-way ANOVA and Tukey's posthoc test. **(b)** Cross-fertilisation experiments with pollen donors and recipients of *Tass4-1* $BC_2F_2$ plants. Fertilisation success rate was calculated for *n*=3-4 crossed ears as the number of grains produced relative to the total number of crossed florets. Crossing *aa bb* donor pollen to an AA BB recipient resulted in significantly reduced fertility success compared to the reciprocal cross ($p<0.05$; two-tailed t-test). **(c)** Number of grains per spike on *Tass4-2* plants. The average number of grains in the three primary spikes was calculated for *n*=2-5 plants. Different letters indicate significant differences at $p < 0.05$ under a one-way ANOVA and Tukey's posthoc test. Note that the statistical analysis includes data in Fig. 8d, which were obtained in the same experiment.

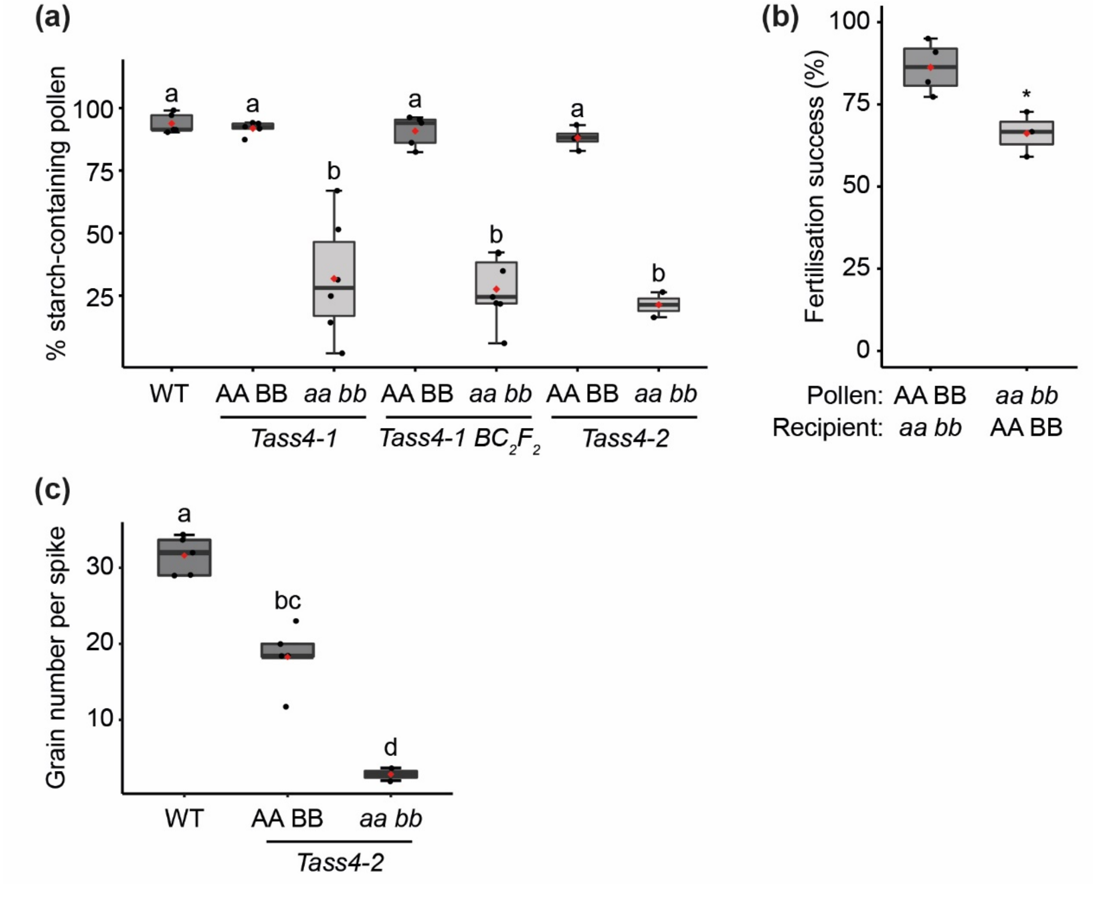

**Table S1** Oligonucleotides used in this study.

| | Forward primer | Reverse primer |
|---|---|---|
| ***Cloning of TaSS4-1B-cTP:pProExHTb*** | | |
| *Ta*SS4-1B-cTP insert | tgagagaagattttcagc | ctgaaaatacaggttttcggtc |
| pProExHTb backbone | ccgaaaacctgtattttcaggctgcggcgcaacgccct | gctgaaaatcttctctcacgctactgcacgggcacgtg |

The 15 bp overlap with the insert for fusion via Gibson assembly is underlined.

| | | |
|---|---|---|
| ***Cloning of TaBGC1-4B-cTP:pProExHTb*** | | |
| *Ta*BGC1-4B-cTP insert | ataggatcctgtgcggcctatatgcc (BamHI) | tatctcgagttatgtgacgatcatcaaattgt (XhoI) |

Restriction enzyme sites used for cloning are underlined. Enzymes are indicated in brackets.

| | | |
|---|---|---|
| ***KASP genotyping*** | | |
| K2166 (*Ta*SS4-1A) | WT: ggctgtattaatactaaaactcac<br>mutant: ggctgtattaatactaaaactcat | common: aacagggaagttatggacaac |
| K2565 (*Ta*SS4-1B) | WT: gtaagatattttcttctccagtaac<br>mutant: gtaagatattttcttctccagtaat | common: agcactcaatagtggaatttcat |
| K1450 (*Ta*SS4-1B) | WT: tggcatgctttgtggtttcag<br>mutant: tggcatgctttgtggtttcaa | common: tccatgggtattgtttcgtca |
| K2244 (*Ta*BGC1-4A) | WT: ttgtcaagagaccatgttcgc<br>mutant: ttgtcaagagaccatgttcgt | common: ctcctagtcgaagtccaacta |
| K3145 (*Ta*BGC1-4B) | WT: tacttttgtctcttcaaattccc<br>mutant: attacttttgtctcttcaaattcct | common: gaagatcttgatgcatctcatc |
| K2275 (*Ta*BGC1-4B) | WT: cttctgtgttacatacatgcag<br>mutant: cttctgtgttacatacatgcaa | common: cttcgagtgtacatgccatc |

The VIC/HEX tail (GAAGGTCGGAGTCAACGGATT) was added to the 5’ end of the wild-type allele-specific primers. The FAM tail (GAAGGTGACCAAGTTCATGCT) was added to the 5’ end of the mutant allele-specific primers.

All primer sequences are given 5’ to 3’.

**Table S2** Codon-optimised coding sequences for *Ta*SS4 and *Ta*BGC1.

| | Coding sequence (5’ to 3’) |
|---|---|
| *TaSS4-1A* | ATGGCGTGTTCCGCGGCAGCTGGCGTAGAGGCCACGGCACTGTTGTCACCCCGTTGTCCTGCACCGTCTCCGCCTGACGGGCG<br>TTCGCGTCGCCGTTTAGCTCTGGCTTCGGGTACTCGCCATCGCAGCCTTCGCGCGGCAGCACAACGCCCTCACAAGAGTGCGA<br>CAGGTGCAGATCCGCTTTATAATAACCGCGCAAACGTCCGTTCCGATGAGGCGTCTGTGAGTGCGGAGAAGGAACGCCAGCGT<br>AACTATAATGATGGGGACGGCATTTCTAATTTGAAATTGGAAGACTTGGTAGGCATGATCCAGAACACAGAAAAAAACATCCT<br>TTTATTGAACCAAGCACGTTTGCAGGCCATGGAACACGCAGATAAAGTCTTGAAAGAGAAAGAAGCCCTTCAGCGTAAGATCA<br>ATATTTTAGAGACGCGCTTAAGCGAAACAGACGAGCAACACAAACTGTCATCTGAGGGGAATTTCTCAGACTCCCCCTTGGCC<br>CTGGAGCTTGGCATCCTGAAGGAAGAAAATATCTTGCTGAAAGAAGACATCGAGTTCTTTAAAACCAAGTTGATCGAGGTGGC<br>CGAGATCGAGGAGGGAATTTTCAAACTGGAGAAGGAGCGTGCGCTGTTGGACGCCTCTCTGCGCGAGCTTGAATCGCGCTTTA<br>TCGCCGCGCAGGCCGATACTATGAAACTTGGTCCCCGCGATGCCTGGTGGGAGAAGGTCGAGAAACTTGAGGATCTTCTTGAA<br>ACCACAGCAAACCAAGTAGAACACGCTGCCGTCATCTTAGACCACAATCATGATTTGCAAGACCGTTTGGATAAGTTAGAAGC<br>TTCATTGCAGGCCGCCAACATCTCGAAATTTTCGTGCTCATTAGTAGACATTCTTCAGCAGAAAGTGAAACTGGTTGAAGAAT<br>GCTTTCAGGCATGCAATTGCGAAATGCATTCGCAGATCGAATTGTACGAACACTCAATTGTCGAATTCCATGACACCCTTAGT<br>AAATTAATCGAAGAATCGGAAAAGCGTAGTCTGGAGAACTTTACTGGAAACATGCCTTCAGAGTTATGGAGCAAGATCTCTCT<br>GTTGACCGACGGCTGGCTTCTTGAAAAAAAAATTAGTTACAATGACGCTTCCATGTTACGCGAGATGGTACATAAGCGTGACT<br>CGCGCCTTCGCGAGGCATATTTAAGTTACCGTGGACGGAAAATCGCGAGGTTATGGACAACTTACTGAAAATGGCCCTGCCA<br>GGTACCTCTAGTGGCCTTCACATCGCCCACATCGCGGCCGAAATGGCGCCCGTGGCGAAAGTGGGGGGCCTTGCAGACGTCAT<br>CTCGGGACTGGGTAAAGCACTGCAGAAGAAAGGACATTTAGTAGAGATCATCCTTCCGAAGTATGATTGCATGCAGGTAGACC<br>AGGTGTCTAATCTGCGCGTATTAGACGTACTGGTCCAAAGCTACTTTGAGGGTAATATGTTTAACAACAAAATCTGGACAGGA<br>ACCGTTGAAGGCTTACCGGTCTATTTTATTGAGCCGCAACACCCGGCGATGTTTTTTTCCCGCGCTCAATACTACGGCGAACA<br>TGATGATTTTAAACGCTTTTCGTATTTCAGCCGCGCAGCCTTGGAACTGTTGTATCAGAGTGGGAAGAAGGTCGATATCATTC<br>ATTGTCATGATTGGCAAACCGCGTTCGTTGCGCCCTTATATTGGGATGTTTATGCTAACTTGGGGTTTAACTCGGCACGTATT<br>TGCTTCACGTGCCATAATTTCGAGTATCAAGGAACAGCGCCTGCCCGTGACTTGGCGTGGTGCGGACTGGATGTTGAACACTT<br>GGATCGCCCGGACCGTATGCGTGACAATTCTCATGGACGTATCAATGCTGTTAAGGGTGCGGTTGTCTATAGTAATATCGTAA<br>CTACTGTGTCCCCGACATATGCCCTGGAGGTACGCTCAGAGGGTGGCGTGGTTTACAAGACACTTTAAAAGTTCATAGTCGC<br>AAGTTCTTGGGCATTCTTAACGGTATCGACACGGACACTTGGAATCCGTCAACAGACCGTTATTTGAAAGTTCAGTATAACGC<br>TAAAGACTTACAAGGTAAGGCAGCGAATAAGGCGGCATTACGCGAGCAGCTGAATCTGGCCTCTGCGTACCCATCGCAGCCGC<br>TTGTTGGTTGTATTACGCGCCTGGTGGCCCAGAAAGGAGTCCACTTAATTCGTCATGCCATCTACAAAACCGCCGAGCTGGGT<br>GGGCAGTTCGTCTTATTGGGGAGTAGTCCGGTACCGGAGATCCAACGTGAGTTCGAAGGAATCGCTGATCACTTTCAAAACAA<br>CAACAACATTCGCTTAATTTTAAAGTACGACGACGCGCTTTCGCACTGCATCTATGCCGCTTCTGACATGTTCATCGTTCCTT<br>CCATCTTCGAGCCATGCGGATTGACTCAAATGATCGCTATGCGCTATGGAAGTGTGCCAATCGTTCGCAAAACAGGTGGTTTA<br>AACGATAGTGTTTTCGACTTTGATGACGAAACGATCCCTATGGAAGTCCGTAACGGATTTACTTTCGTAAAAGCCGACGAGCA<br>AGGGTTGTCTTCAGCTATGGAACGCGCGTTTAATTGCTATACCCGCAAACCGGAAGTTTGGAAACAGCTGGTTCAAAAGGACA<br>TGACTATTGATTTTTCCTGGGATACAAGTGCGTCCCAGTATGAGGACATCTACCAGAAAGCAGTGGCCCGCGCGCGCGCTGTT<br>GCA |
| *TaSS4-1B* | ATGGCTTGCTCTGCAGCCGCTGGGGTAGAGGCAACAGCTCTGCTGAGTCCTCGCTGTCCAGCACCGTCGCCTCCGGACGGACG<br>CTCACGCCGTCGCCTGGCCTTGGCTTCGCGTACACGTCATCGTAGTTTGCGTGCTGCGGCGCAACGCCCTCATAAGTCCACAA<br>CAGGAGCTGACCCCCTTAATAACCGTGCGAACGTCCGTAGTGATGAAGCCGCCGTGTCGGCTGAAAAGGAGCGCCAACGTAAG<br>TATAATGACGGAGATGGTATCTCAAACTTACAGCTTGAGGATCTTGTCGGCATGATCCAGAACACCGAAAAGAACATCCTGTT<br>GCTTAACCAGGCGCGCCTGCAGGCGATGGAGCATGCGGACAAGATCCTGAAAGAAAAGGAGGCACTTCAACGTAAGATTAACA<br>TCTTAGAAACCCGTTTAAGCGAGATTGACTCCCAACACAAGTTGTCATCGGAGGGAAACTTCTCGGACTCGCCTCTGGCACTT<br>GAGTTTGACGTGCTTAAGGAAGAGAACATCGTATTGAAGGAAGACATTGAATTTTTCAAGACCAAGTTAATCGAAGTAGCTGA<br>GACTGAGGAGGGTATCTTTAAGCTTGAAAAGGAGCGTGCCCTTCTGGACGCGTCCTTGCGCGAGTTGGAGTCCCGCTTCATCG<br>CCGCTCAAGCTAACATGATGAAATTAGGGCCACGTGACGCATGGTGGGAGAAGGTGGAAAAATTGGAGGACTTATTAGAAACC<br>ACAGCTAATCAGGTAGAACATGCTGCGGTGATCCTTGACCGTAATCACGACCTGCAAGACCGTTTAGAAAAACTGGAGGCTTC<br>TCTGCAGGCCGCTAATATCAGCAAGTTCTCGTGTTCATTAGTAGACTTGCTTCAGCAGAAAGTAAAATTAGTGGAGGAGCGCT<br>TCCAAGCATGCAACCGCGAAATGCATTCTCAAATTGAATTGTATGAGCATTCGATTGTTGAATTTCACGATACATTATCAAAG<br>CTGATCGAAGAATCCGAAAAGCGTTCACTGGAGAATTTTACAGGGAATATGCCATCGGAGCTTTGGAGCAAAATTTCATTACT<br>GACCGATGGCTGGCTGTTGGAGAAAAAGATCAGTTACAGCGATGCCAGCATGTTACGCGAAATGGTCCAGAAACGCGACAATC<br>GTTTGCGCGAAGCATACTTAAGTTATCGTGGACAGAAAACCGCGAAGTCATGGATAATTTGCTTAAAATGGCTCTGCCGGGA<br>ACTTCTAGCGGGTTACACATTGCTCATATTGCGGCGGAGATGGCGCCGGTTGCCAAGGTGGGCGGCCTTGCGGACGTAATTTC<br>TGGATTAGGCAAAGCATTGCAAAAAAAGGGCCACTTGGTAGAGATCATTCTGCCGAAATATGACTGTATGCAAGTGGATCAGG<br>TTTCCAATCTGCGCGTCTTAGATGTATTAGTCCAGTCCTACTTCGAAGGCAATATGTTTAATAACAAGATCTGGACAGGCACC<br>GTAGAAGGACTGCCAGTTTACTTCATCGAACCGCAGCACCCTGCTATGTTCTTCAGCCGCGCTCACTATTACGGGGAACACGA<br>TGATTTTAAGCGTTTTTCGTATTTCAGTCGCGCTGCACTGGAATTACTGTACCAGTCAGGCAAAAAGGTTGACATCATCCATT<br>GCCATGACTGGCAGACTGCCTTTGTAGCCCCTTTGTATTGGGATGTGTATGCGAACCTGGGATTTAATTCCGCTCGTATCTGT<br>TTTACATGTCACAATTTTGAATACCAAGGCACGGCGCCCGCTCGTGACTTAGCATGGTGCGGCCTTGATGTCGAACACTTAGA<br>TCGCCCGGATCGTATGCGCGATAATTCACACGGCCGCATCAATGCAGTGAAGGGTGCGGTAGTTTATAGTAACATCGTTACAA<br>CGGTCTCGCCAACTTACGCTCTGGAGGTCCGTAGCGAAGGAGGACGCGGGTTGCAAGATACGCTGAAAGTACACTCTCGTAAG<br>TTTTTGGGTATCTTGAATGGGATTGATACAGATACTTGGAACCCGTGCACCGATCGTTATCTTAAGGTCCAATATAACGCCAA<br>AGACTTACAAGGGAAAGCGGCTAATAAAGCTGCCCTTCGTGAACAGCTTAACCTGGCGTCCGCGTACCCCAGTCAGCCGCTTG<br>TTGGGTGTATCACGCGCTTGGTCGCACAGAAGGGTGTGCACCTGATCCGCCACGCAATCTATAAGACAGCCGAACTGGGTGGG<br>CAATTCGTGTTGCTGGGCTCAAGCCCAGTGCCCGAGATCCAACGCGAATTCGAGGGGATTGCGGACCACTTCCAAAACAACAA<br>TAACATTCGTTTAATCCTTAAGTATGATGACGCCCTTAGCCATTGCATTTATGCGGCATCTGATATGTTCATTGTGCCGTCTA<br>TCTTTGAGCCGTGTGGACTTACACAGATGATCGCTATGCGTTATGGTAGTGTTCCCATTGTGCGTAAAACTGGTGGATTGAAC<br>GACAGTGTTTTTGACTTCGACGATGAGACAATCCCTATGGAAGTGCGTAACGGCTTCACATTTGTGAAAGCTGATGAGCAAGG<br>TTTGTCCAGTGCTATGGAACGCGCATTTAACTGCTATACCCGCAAGCCAGAGGTCTGGAAGCAATTGGTGCAGAAAGACATGA |

| | |
|---|---|
| | CTATTGATTTCTCCTGGGACACGTCTGCCTCCCAGTACGAGGATATTTATCAAAAAGCAGTAGCACGTGCCCGTGCAGTAGCG |
| *TaBGC1-4A* | ATGCCGCCGTTTCTTCCCAGCTTACCCCTGCCCGCACTTACTCTTTCATTACCATTACCGCCGGCGCCGGCACCACGCCGTCA<br>TCGTGTATTTGCTGCTGCCGCGGCATATGGCCCACAACCCTGTCGCGGACGTGTGTGTGTGTGTGCAGCTTACCGCCCTCCTC<br>CGCGTCAACCATATCGCCGTCAACCCGCGCCCGCTCCTGCCCCTCGCCCGCCCAATGCGCCTGCTCCACCACAACGTGGTCCG<br>CGCGGCCAGGAGGAACTGGAAGAGGCAATCTATGATTTTATGCGCCGTTCCGATAAACCAGGAGCCTTCCCCACGCGCGCTGA<br>ACTGCTTGCGGCAGGGCGTGCAGACCTTGCGGCCGCGGTTGAGAGTTCAGGAGGTTGGTTATCTCTGGGGTGGTCATGGAGTA<br>GTGATGATGATGCGCGCCGTCCAGCAGCAAGTACTGCCGGTCCTGGGGTCCACCCGGAATACCCCCCTGAAGCTGGCCCAAGT<br>GGCCGCCCACCTAATAGCGCGGCCGATTCTGTGCGCGAACAGCAAGAGCCGGCTCCATCCGGCCGCCAGCCAGAAACGGAGGA<br>GACCGAAGAGGCGGGATCGGGTGCAGGGTTAGAGGGTATGCTGGCCCGCTTGCGCCGTGAGCGCGAACGCGCTCGCCCACCTC<br>CCCGCAGCAAGAACCAGGCCGGCGGTCGCGGCCAGAACGGAGCCCTTATGAACCATAACGGCGCTCCCTCCCGCAGTCCAACT<br>AATGGCATGTACACACGCCGTATTCCCGTTAACGGGAACATCCATCGTAGTCACTCTCAGAATGGTATTCCCGAAGATAACAA<br>GTCGTCGTCTTCCGCTAATGATGCGTGGCGCACGTGGAGCCTTGACAAGAGCCGCTTTTCGGACTTTGAAGCAGCTGAAATCC<br>ATCCGCTTAGTCGTAAGCCACCGAAGCACGTTGATTTGAACACAGTTCTGATTGAGGATGACGTTCCAGGACCCTCCAATGGG<br>GTGGTAATTAACGATTACCCGAGCGACCACGTAGATTCGGAGCGCGACGAGATTCACGCTCGCTTCCAAAACCTGGAATTCGA<br>CTTGGCGGACTCTCTTAAGACCTTGCGCTCGCGTTTCGATGGGGTATCATCATATATGTCAAACGGAGAGGAGGCGGATGTAG<br>TCAATGGGTTTTCAGACGATTGGGAATTCGAAGAAACGAAAGTCATGCACGCACAAGAGGAATTGCGTACAATCCGCGCAAAG<br>ATCGCGGTGTTGGAGGGGAAGGTAGCACTGGAAATTATCGAAAAAAACAAAATTATCGAGGAGAAGCAGACCCGCTTAGACGA<br>GGTTGAAAAAGCCCTGTCTGAGCTGCGTACTGTTTCAGTAGTATGGCCTAACCCAGCATCCGAAGTGTTATTAACTGGATCCT<br>TCGACGGCTGGACGTCACAACGCCGCATGGAACAGAGCGAGTCGGGAATCTTCAGCTACAATCTTCGTCTTTACCCCGGACGT<br>TATGAGATTAAGTTTATCGTCGATGGAGTGTGGAAAAACGATCCACTTCGCCCAAGTGTGAACAACCACGGTAATGAGAACAA<br>CCTTATGATCGTCACT |
| *TaBGC1-4B* | ATGCCTCCTTTCCTGCCCAGTTTACCATTACCCGCGCTGACTTTGCCGTTGCCCTTACCACCATTGTTAACCGCTCCCGCACC<br>CCGTCGCCACCGTGTGTTCGCCGCAACAGCCGACGGACCTCAACCTTGCCGCGGCCGTGTGTGCGTATGTGCGGCCTATATGC<br>CCCCGCCGCGTCAGCCTTACCGTCGCCCAGCGCCGGCACCGGCTCCTGCCCCGCGTCCTAGCAACGCTCCGGCTCCGGCCCCA<br>CCACAACGTGGACCACGCGACCAAGAGGAGTTAGAAGCAGCTATTTACGACTTCATGCGCCGCTCTGATAAGCCCGGGGCCTT<br>TCCGACACGTGCTGAGCTGCTTGCAGCTGGACGCGCAGATTTAGCAGCTGCAGTTGAGAGCTCAGGTGGCTGGTTATCGCTGG<br>GCTGGTCTTGGTCTTCTGATGACGACGCACGCCGCCCAGCTGCATCTACGGCCGGTCCAGGGGTGCACCCTGATTATCCGCCG<br>GAGGCAGGGGCAAGTGGGCGTGCCCCTAATGCAACAGCGGACTCTGTGCGCGAGCAGCAGGAGCCAACACCAAGCGGACGCCA<br>ACCAGAAACGGAGGAGACGCAAGAAGCAGGGTCTGGGGCTGGTTTAGAGGGGATGTTGACCCGCTTACGCCGTGAGCGTGAAC<br>GCGCTCGCCCACCGCCTCGCAGTAAGAATCGCGCGGGGGGCCAAGGACAGAACGGTGCCTTGATGAACCACAATGGGGCGCCT<br>TCGCGCTCTCCAACTGATGGTATGTATACCCGTCGCATCCCAGTCAACGGGAACATCCACCGTTCTCACTCACAAAATGGAAT<br>TCCAGAGGATAACAAAAGTTCATCTTCTGCCAACGACGCTTGGCGCACCTGGTCCTTAGACAAGAGCCGTTTTTCGGATTTCG<br>AAGCGGCCGAGATTCATCCGTTAAGTCGTAAGCCGCCGAAACGTGCCGATCTTGACACCGTTTTGATCGAAGACGATGTACCC<br>GGACCCTCCAATGGGGTAGTAATCAACGATTATCCATCTGACCACGTTGACTCTGAACGTGACGAGATTCATGCCCGCTTTCA<br>AAACCTGGAATTCGACTTAGCTGACTCCCTTAAAACGTTACGCTCCCGCTTTGATGGCGTTAGCTCATATATGAGCAACGGGG<br>AGGAAGCAGATGTCGTTAACGGCTTTTCGGACGATTGGGAATTCGAAGAGACTAAGGTTATGCACGCCCAAGAAGAACTGCGT<br>ACAATCCGCGCTAAAATTGCTGTTTTAGAGGGAAAGGTTGCTTTGGAGATCATTGACAAAAACAAAATCATTGAAGAGAAACA<br>GACCCGTTTAGACGAAGTTGAAAAAGCGTTGAGTGAATTACGCACAGTGAGCGTAGTGTGGCCGAATCCCGCTTCGGAAGTTC<br>TTCTGACAGGATCTTTTGACGGTTGGACGTCCCAACGTCGTATGGAGCAGTCTGAGGGGGGGATTTTCAGCTATAATTTACGC<br>TTATACCCAGGCCGCTACGAAATTAAATTCATCGTGGACGGCGTTTGGAAAAACGATCCGCTTCGCCCAACCGTCAACAATAA<br>TGGAAATGAAAACAATTTGATGATCGTCACA |
